# Phosphate-induced resistance to pathogen infection in Arabidopsis

**DOI:** 10.1101/2021.12.09.471930

**Authors:** Beatriz Val-Torregrosa, Mireia Bundó, Héctor Martín-Cardoso, Marcel Bach-Pages, Tzyy-Jen Chiou, Victor Flors, Blanca San Segundo

## Abstract

In nature, plants are concurrently exposed to a number of abiotic and biotic stresses. Our understanding of convergence points between responses to combined biotic/abiotic stress pathways remains, however, rudimentary. Here we show that *MIR399* overexpression, loss-of-function of *PHO2* (*PHOSPHATE2*), or treatment with high Pi, is accompanied by an increase in phosphate (Pi) content and accumulation of reactive oxygen species (ROS) in *Arabidopsis thaliana*. High Pi plants (e.g. miR399 overexpressor, *pho2* mutant, and plants grown under high Pi supply) exhibited resistance to infection by necrotrophic and hemibiotrophic fungal pathogens. In the absence of pathogen infection, the expression level of genes in the salicylic acid (SA)- and jasmonic acid (JA)-dependent signaling pathways was higher in high Pi plants compared to wild type plants, which is consistent with increased levels of SA and JA in non-infected high Pi plants. During infection, an opposite regulation in the two branches of the JA pathway (ERF1/PDF1.2 and MYC2/VSP2) occurs in high Pi plants. Thus, while the ERF1-PDF1 branch positively responds to fungal infection, the MYC2/VSP2 branch is negatively regulated during pathogen infection in high Pi plants. This study supports that Pi accumulation promotes resistance to infection by fungal pathogens in Arabidopsis, while providing a basis to better understand crosstalk between Pi signaling and hormonal signalling pathways for modulation of plant immune responses.

**Significance statement:** This study highlights the importance of phosphate (Pi) in regulating immune responses, hence, disease resistance in *Arabidopsis thaliana*. Increasing Pi content either by *MIR399* overexpression (or loss-of-function of *PHOSPHATE2*), as well as by Pi treatment enhances resistance to infection by necrotrophic and hemibiotrophic fungal pathogens through modulation of SA- and JA-dependent signaling pathways. These results also support that miR399 functions as a regulator of Arabidopsis immunity.

## INTRODUCTION

In nature, plants are simultaneously exposed to a combination of biotic and abiotic stresses that are diverse in time and space, which requires proper integration and crosstalk between different stress response pathways. Exposure to a single stress might, however, impact the plant response to another stress (Kissoudis *et al.*, 2014; Nejat and Mantri, 2017; Pandey *et al.*, 2017). For instance, plant immune responses and disease resistance can be altered in plants exposed to drought or high salinity (Yasuda *et al.*, 2008; Atkinson and Urwin, 2012; Bostock *et al.*, 2014). Inappropriate supply of mineral nutrients (e.g. nitrogen supply) might also impact disease severity (Snoeijers *et al.*, 2000; Ballini *et al.*, 2013). However, the effect of combined abiotic and biotic stress factors might vary depending on the nature of these interactions, and the plant response to simultaneously or sequentially applied stresses cannot be simply inferred from responses to individual stresses (Prasch and Sonnewald, 2013; Coolen *et al.*, 2016; Pandey *et al.*, 2017; Nobori and Tsuda, 2019). As stress responses are costly, when facing with multiple stresses simultaneously, plants need to prioritize their stress responses for efficient use of finite resources, in accordance with the optimal defense theory (ODT) (Meldau *et al.*, 2012). According to ODT, stress responses are prioritized in the most valuable parts (Keith and Mitchell-Olds, 2017), and recent findings indicate that Arabidopsis plants spatially separate contrasting stress responses in leaves of different ages (e.g. young leaves exhibit higher biotic stress responses but lower abiotic stress responses compared with old leaves) (Berens *et al.*, 2019; Wolinska and Berens, 2019). To date, little information is available on the molecular mechanisms by which biotic and abiotic stress responses are differentially prioritized in plants, and how they adapt to conflicting stresses for optimal responses.

To defend themselves against pathogens, plants have evolved an innate immune system in which many interconnected processes are involved (Jones and Dangl, 2006). Pathogen-induced pathways are defined principally according to the molecules recognized by the host plant (Jones and Dangl, 2006; Boller and Felix, 2009; Thomma *et al.*, 2011; Couto and Zipfel, 2016; Upson *et al.*, 2018). Plants recognize pathogen-associated molecular patterns (PAMPs) by receptors at the plasma membrane, which triggers the induction of multiple cascades leading to the induction of immune responses, referred to as PAMP-triggered immunity (PTI). Components of PTI include the reinforcement of cell walls, accumulation of reactive oxygen species (ROS), activation of phosphorylation cascades, production of antimicrobial compounds and accumulation of pathogenesis-related (PR) proteins, among others (Andersen *et al.*, 2018). Some successful pathogens can overcome PTI by delivering effectors into plant cells that suppress PTI, thus, leading to disease susceptibility. In turn, some plants have evolved another immune response in which microbial effectors (or host proteins modified by effectors) are recognized by intracellular receptor proteins encoded by resistance (*R*) genes (Han, 2019). This recognition triggers a rapid and robust defense response, the so called effector-triggered immunity (ETI) (Jones and Dangl, 2006). ETI is often accompanied by a hypersensitive response (HR) at the infection site, a form of programmed cell death (Thakur *et al.*, 2019). However, some PAMPs (e.g the bacterial Harpin HrpZ protein) can also induce HR in plants (Chang and Nick, 2012).

Among ROS, H_2_O_2_ is relatively stable and is an important molecule regulating plant immunity (Torres *et al.*, 2006). H_2_O_2_ might have a direct antimicrobial role against the invading pathogen and also provokes cross-linking of cell wall components to arrest pathogen invasion. ROS also function as molecules for the activation of defense mechanisms, and triggers localized cell death around the infection site (Torres *et al.*, 2002). Phytohormones, together with ROS, provide important signals to help orchestrate plant responses to abiotic and biotic stresses. On the other hand, immune responses are largely coordinated by the phytohormones salicylic acid (SA), ethylene (ET), and jasmonic acid (JA) (Aerts *et al.*, 2021). Plant hormones do not function independently, but synergistic or antagonistic interactions between hormone pathways ultimately drive the fine-tuning of plant defense responses. At the organismal level, hormone crosstalk might also balance trade-offs between conflicting biotic and abiotic stress responses (i.e. prioritization of responses in leaves of Arabidopsis plants) (Berens *et al.*, 2019; Wolinska and Berens, 2019). Sugar and ROS have also been proposed as candidates signaling molecules to regulate prioritization between biotic and abiotic stress responses (Wolinska and Berens, 2019).

Plant miRNAs are a class of small RNA molecules that mediate post-transcriptional gene silencing through sequence complementarity with cognate target transcripts (Bartel, 2004; Xie *et al.*, 2005). They are transcribed from *MIR* genes as long precursor transcripts that adopt a stem-loop structure by self-complementarity that are processed by a RNase III DICER-like, typically DCL1, to produce a double stranded duplex, the miRNA-5p/miRNA-3p duplex (previously named as miRNA/miRNA*) (Borges and Martienssen, 2015). The functional strand of the duplex is selectively loaded into an ARGONAUTE (AGO) protein, the effector protein of the RISC (RNA-induced silencing complex), and guide post-transcriptional gene silencing by cleaving the target transcripts or by translational inhibition (Llave *et al.*, 2002; Brodersen *et al.*, 2008). The important role of plant miRNAs in diverse developmental processes and adaptation to environmental stresses is well documented (Chen, 2009; Seo *et al.*, 2013; Staiger *et al.*, 2013; Song *et al.*, 2019).

The first plant miRNA demonstrated to be involved in immunity was the Arabidopsis miR393. Here, perception of the elicitor flg22 induces miR393 accumulation and down regulation of auxin receptors, resulting in resistance to bacterial pathogens (Navarro *et al.*, 2006). Since then, other miRNAs controlling diverse processes have been shown to be involved in Arabidopsis immunity, either ETI or PTI (Jagadeeswaran *et al.*, 2009; Shivaprasad *et al.*, 2012; Seo *et al.*, 2013; Staiger *et al.*, 2013; Huang *et al.*, 2016). Some examples are: miR160, miR396, miR398, miR400, miR472, miR773, miR844, miR858, miR863, miR156 (Li *et al.*, 2010; Boccara *et al.*, 2014; Park *et al.*, 2014; Lee *et al.*, 2015; Niu *et al.*, 2016; Camargo-Ramírez *et al.*, 2017; Salvador-Guirao *et al.*, 2018; Yin *et al.*, 2019). Depending on their target gene, these miRNAs can function as positive or negative regulators in fine-tuning immune responses. Even though pathogen infection has been shown to induce alterations in the expression of a plethora of Arabidopsis miRNAs, the mechanistic role of these miRNAs in processes underling immunity is often unclear. Additionally, most studies on miRNAs involved in plant immunity have been carried out in the interaction of Arabidopsis plants with the bacterial pathogen *Pseudomonas syringae*, and less is known on miRNAs involved in resistance against fungal pathogens.

On the other hand, distinct miRNAs have been associated with regulation of nutrient homeostasis in plants (Paul *et al.*, 2015). Perhaps the best known example is miR399 which is involved in the control of phosphate (Pi) homeostasis in Arabidopsis plants (Chiou *et al.*, 2006). Under limiting Pi conditions, miR399 accumulation increases and causes repression of its target gene, *PHO2* (*PHOSPHATE2*) encoding an E2 ubiquitin-conjugating enzyme responsible of degradation of phosphate transporters (Fujii *et al.*, 2005; Chiou *et al.*, 2006; Liu *et al.*, 2012; Huang *et al.*, 2013; Kraft *et al.*, 2016). Hence, miR399 accumulation in response to Pi starvation relieves negative post-transcriptional control of Pi transporters and promotes uptake of Pi in Arabidopsis plants. The miR399/PHO2 regulatory module has been also recognized as a regulator of Pi homeostasis in rice (Chien *et al.*, 2017; Puga *et al.*, 2017).

In this work, we investigated whether miR399 plays a role in disease resistance in Arabidopsis. We show that miR399 overexpression and loss-of-function of *pho2* causes an increase in Pi level, these plants also exhibiting resistance to infection by necrotrophic (*Plectosphaerella cucumerina*) and hemibiotrophic (*Colletotrichum higginsianum*) fungal pathogens. Growing Arabidopsis plants under high Pi supply also increases resistance to infection by these pathogens. In the absence of pathogen infection, plants that overaccumulate Pi (e.g. miR399 overexpressor, *pho2* mutant, wild type plants grown under high Pi supply) showed ROS accumulation, increased SA and JA levels, and up-regulation of genes involved in SA- and JA-dependent defense pathways. Pathogen infection was found to be associated with a higher production of ROS in high Pi plants. To note, during pathogen infection, induction of the ERF1 branch of the JA pathway, but repression of the MYC2/VSP2 branch, occurs in high Pi plants. Overall, our results support that an increase in Pi content has an impact on hormone networks regulating Arabidopsis defense and promotes resistance to pathogen infection in Arabidopsis. These results are markedly different from those recently reported on rice (Campos-Soriano *et al.*, 2020) where miR399 overexpression and high Pi supply were found to enhance susceptibility to infection by the rice blast fungus *Magnaporthe oryzae*. These findings illustrate the need of investigating the effects of nutrient supply on the expression of immune responses and disease resistance on a case-by-case basis.

## RESULTS

### Resistance to infection by fungal pathogens in Arabidopsis plants overexpressing *MIR399*

In Arabidopsis, the miR399 family comprises 6 members, *MIR399a-f.* Of them, miR399f has identical mature sequences in Arabidopsis and rice. Previous studies indicated that transgenic Arabidopsis and rice plants overexpressing miR399f overaccumulate Pi in leaves (Chiou *et al.*, 2006; Campos-Soriano *et al.*, 2020). In this work, we investigated whether miR399f overexpression, and subsequent Pi accumulation, has an effect on resistance to pathogen infection in Arabidopsis. Towards this end, transgenic plants overexpressing *MIR399f* (hereinafter referred to as miR399 OE plants) were generated. Compared to wild type plants, miR399 OE lines accumulated precursor and mature miR399 sequences, which was accompanied by a decrease in *PHO2* transcript level, and overaccumulation of Pi in rosette leaves (Figure 1a).

**Figure 1.**
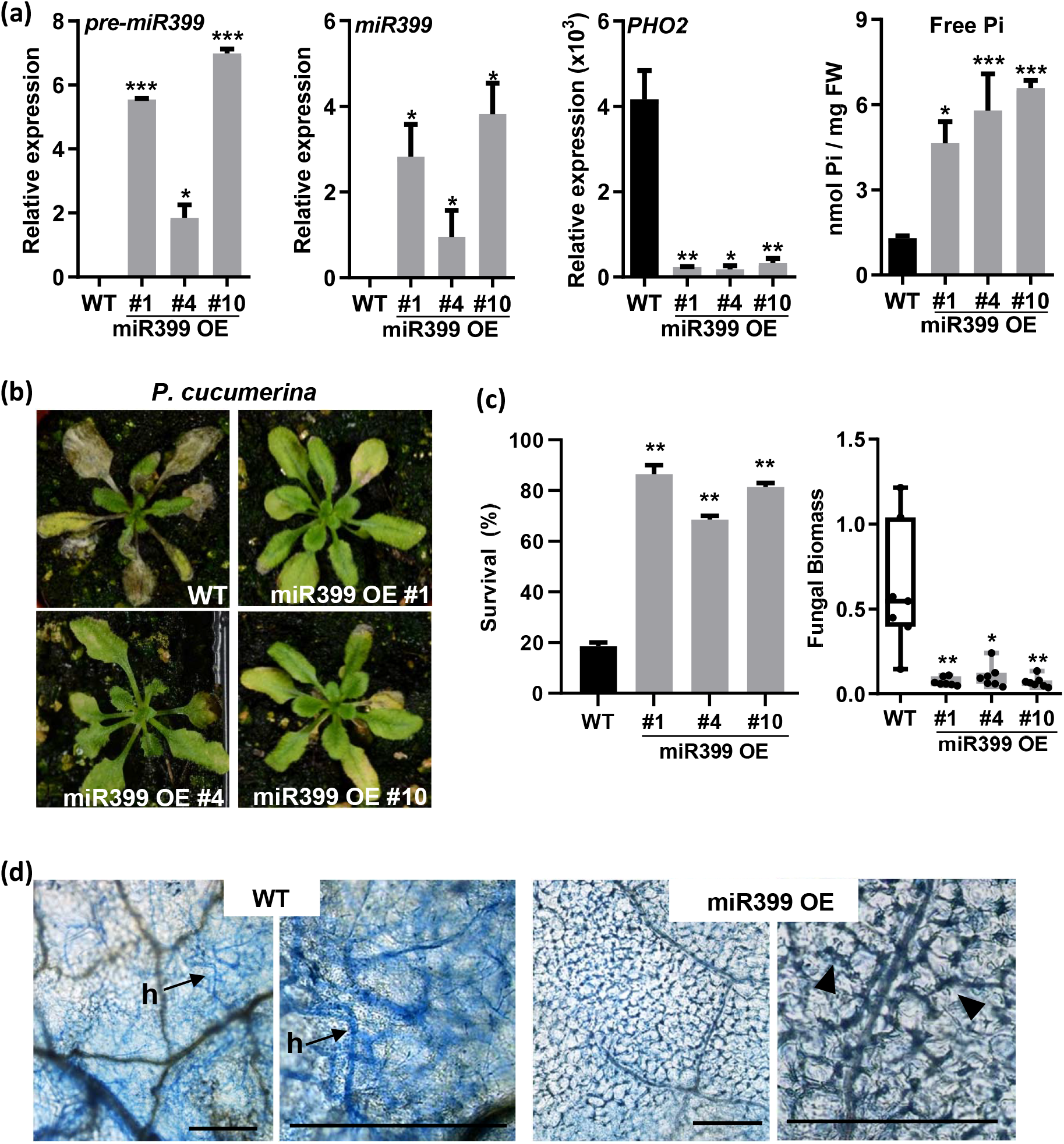
Resistance of miR399 OE plants to infection by the necrotrophic fungus *P. cucumerina.* Homozygous miR399 OE lines (#1, #4 and #10) and wild type (WT) plants were grown in soil for three weeks and then assayed for disease resistance. Three independent experiments were carried out with at least 12 plants/line each experiment. (a) Accumulation of precursor (pre-miR399) and mature miR399 sequences were determined by RT-qPCR and stem-loop RT-qPCR, respectively. Expression of the miR399 target *AtPHO2* by RT-qPCR. The Arabidopsis *β-tubulin2* gene (At5g05620) was used to normalize transcript levels (relative expression). The accumulation of free Pi in leaves is shown (right panel). Bars represent mean ± *SEM* of 3 biological replicates with at least three plants per replicate (*t* test, \**p* ≤ 0.05; \*\**p* ≤ 0.01; \*\*\**p* ≤ 0.001). (b) Plants were spray-inoculated with *P. cucumerina* spores (5 x 10 spores/ml). Pictures were taken at 7 days post inoculation (dpi). (c) Survival ratio of WT and miR399 OE plants at 7dpi. Quantification of *P. cucumerina* DNA was performed by qPCR using specific primers of *P. cucumerina β-tubulin* at 7 dpi. Values of fungal DNA were normalized against the Arabidopsis *UBIQUITIN21* gene (At5g25760). Comparisons have been made relative to WT plants. Data are mean± *SEM* (*n* 7) (*t* test, *\*p≤* 0.05, *\*\*p* ≤ 0.01). (d) Trypan blue staining of *P. cucumerina*-infected leaves of wild type and miR399 OE plants (7 dpi). h, hyphae. Arrows and arrowheads indicate fungal hyphae and death cells, respectively. Higher magnifications are shown (WT and miR399 OE, right panels). Bars represent 300 μm.

The miR399 OE lines were challenged with the fungus *P. cucumerina*, the causal agent of the sudden death and blight disease in many dicotyledonous species. The Arabidopsis/*P. cucumerina* pathosystem emerged as the model system for studies on disease resistance against necrotrophic fungi (Ton and Mauch-Mani, 2004; Sánchez-Vallet *et al.*, 2012). At the time of pathogen inoculation (3-week-old plants), no obvious phenotypic differences were observed between miR399 OE plants and wild type plants (Figure S1a, b). However, at a later developmental stage, the miR399 and *pho2* plants displayed symptoms of Pi excess (e.g. chlorosis on mature leaves of adult plants), most probably, because of Pi accumulation overtime (results not shown; similar results were previously reported by Chiou *et al.*, 2006).

Upon pathogen challenge, wild type plants were severely affected by *P. cucumerina*, while the miR399 OE plants consistently exhibited enhanced resistance (Figure 1b). While 75-85% of miR399 OE plants survived at 7 dpi, only 20% of the wild type plants were able to overcome infection (Figure 1c, left panel). Quantitative PCR (qPCR) measurement of fungal DNA confirmed less fungal biomass in leaves of miR399 OE plants compared with wild type plants infected with *P. cucumerina* (Figure 1c, right panel), which is consistent with the phenotype of resistance that is observed in miR399 OE plants.

Trypan blue staining was used to visualize both fungal structures and dead cells in the fungal-infected leaves of wild type and miR399 OE plants. Whereas extensive fungal growth occurred in leaves of wild type plants, no fungal growth could be observed in leaves of miR399 OE plants (Figure 1d). Instead, scattered groups of dead cells were visualized in *P. cucumerina*-infected miR399 OE plants (Figure 1d, right panels).

Since *P. cucumerina* is a necrotrophic fungus, we hypothesized that the effect of miR399 overexpression in disease resistance might be dependent on the lifestyle of this pathogen. Accordingly, we investigated resistance of miR399 OE plants to infection by the hemibiotrophic fungus *Colletotrichum higginsianum*. This fungus causes the anthracnose leaf spot disease on *Brassica* species, including *A. thaliana* (O’Connell *et al.*, 2004). The miR399 OE plants showed resistance to *C. higginsianum* infection relative to wild type plants, as revealed by quantification of diseased leaf area and fungal biomass (Figure 2a). As observed in infection assays with *P. cucumerina*, the *C. higginsianum-infected* miR399 OE plants exhibited groups of dead cells in their leaves (Figure 2b).

**Figure 2.**
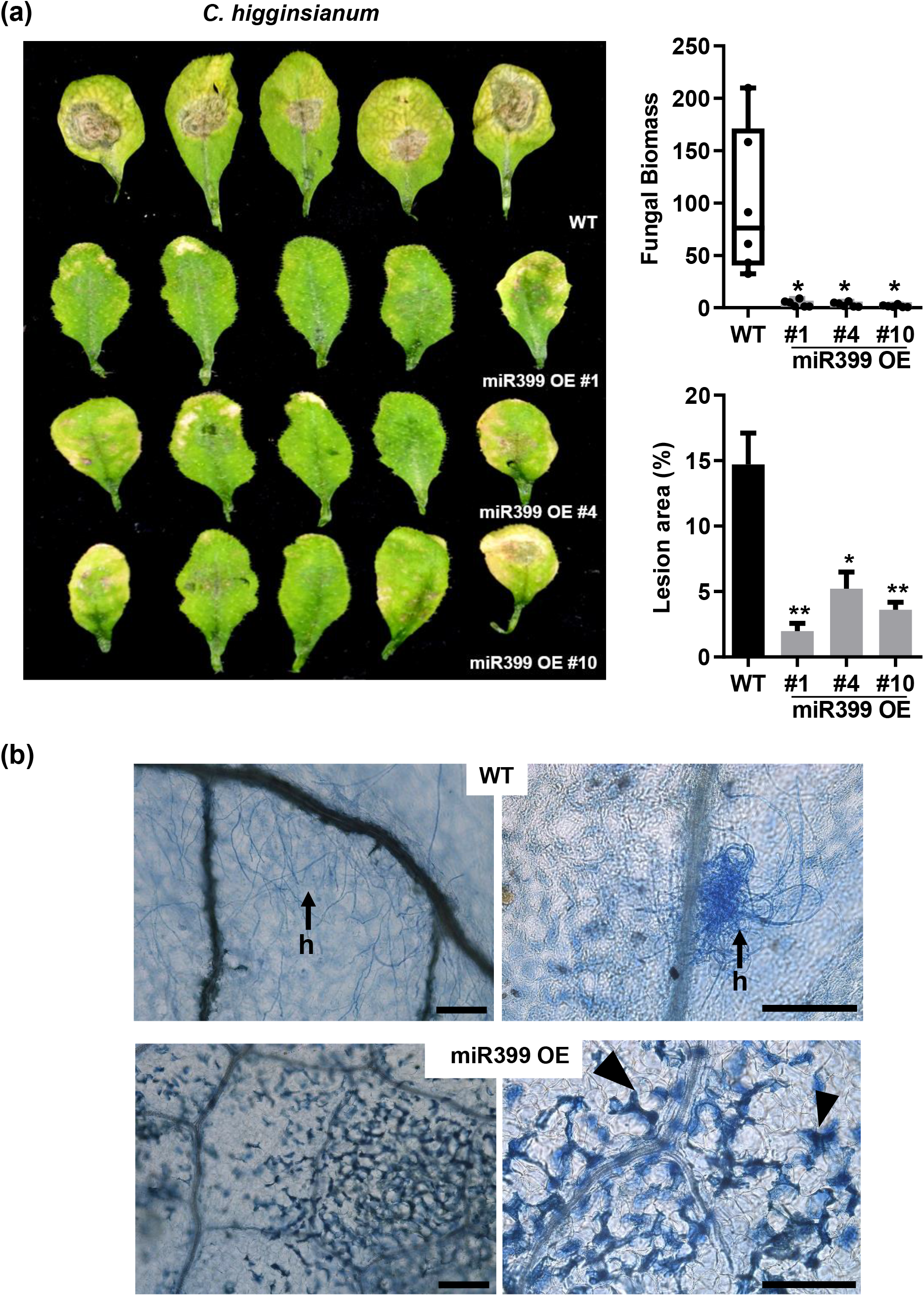
Resistance of miR399 OE plants to infection by the hemibiotrophic fungus *C. higginsianum*. Leaves were locally inoculated with a spore suspension at 4 x 10^6^ spores/ml. Results are from one out of three independent experiments performed with three independent miR399 OE lines (lines 1, 4 and 10) and wild type plants which gave similar results. At least 12 plants/genotype were assayed in each experiment. (a) Disease symptoms at 7 days after inoculation with fungal spores. Diseased leaf area was quantified using image analysis software (ImageJ). Quantification of *C. higginsianum* DNA was carried out by qPCR using specific primers for the *C. higginsianum ITS2* (*Internally transcribed spacer 2*) gene at 7 days post inoculation (dpi). Values are fungal DNA levels normalized against the Arabidopsis *UBIQUITIN21* gene (At5g25760). Comparisons have been made relative to WT plants. Histograms show the mean ± *SEM (t* test, \**p* ≤ 0.05; \*\**p* ≤ 0.01). (b) Trypan blue staining of *C. higginsianum-infected* leaves of wild type and miR399 OE plants at 7 dpi. h, hyphae. Arrows and arrowheads indicate fungal hyphae and cell death, respectively. Higher magnification of these regions are shown (right panels). Bars represent 100 μm.

Collectively, these results demonstrate that miR399 overexpression enhances resistance to infection by necrotrophic (*P. cucumerina*) and hemibiotrophic (*C. higginsianum*) fungal pathogens. Hence, it is likely that miR399-mediated resistance in Arabidopsis does not depend on the lifestyle of the fungus. A pattern of cell death occurs in the fungal-infected miR399 OE plants.

### *pho2* mutant plants exhibit resistance to infection by fungal pathogens

miR399 targets and suppresses *PHO2* expression, this gene encoding the ubiquitin conjugating enzyme gene that mediates the degradation of Pi transporter proteins (Fujii *et al.*, 2005; Chiou *et al.*, 2006; Huang *et al.*, 2013). A loss-of-function allele of *PHO2* was previously described (Delhaize and Randall, 1995; Aung *et al.*, 2006). This mutant allele possesses a single nucleotide mutation that causes premature termination and loss of ubiquitin conjugation activity of PHO2 (Figure S2a). *pho2* resembles miR399 overexpressing plants in that they both show Pi overaccumulation in leaves resulting from increased Pi uptake from roots and root-to-shoot translocation (Aung *et al.*, 2006) (Figure S2b). We therefore hypothesized that loss-of-function of *PHO2* might result in similar disease phenotype as suppression of *PHO2* by overexpression of miR399.

The *pho2* plants were then examined for pathogen resistance. No phenotypic differences were observed between *pho2* and wild type plants at the time of inoculation (Figure S2c). Consistent with the disease phenotype observed in miR399 OE plants, the *pho2* mutant exhibited resistance to infection by *P. cucumerina* (Figure 3a), which was confirmed by quantifying survival ratio of the infected plants and the amount of fungal biomass (Figure 3b). Trypan blue staining revealed a pattern of dead cells in leaves of *pho2* mutants that have been inoculated with *P. cucumerina* spores (Figure 3c), as it was also observed in miR399 OE plants. Finally, the *pho2* plants also showed resistance to *C. higginssianum* infection as compared with wild type plants (Figure 3d). Taken together, results here presented support that both *pho2* and miR399 overexpressing plants accumulate Pi in leaves and exhibit resistance to infection by fungal pathogens with a necrotrophic or hemibiotrophic lifestyle.

**Figure 3.**
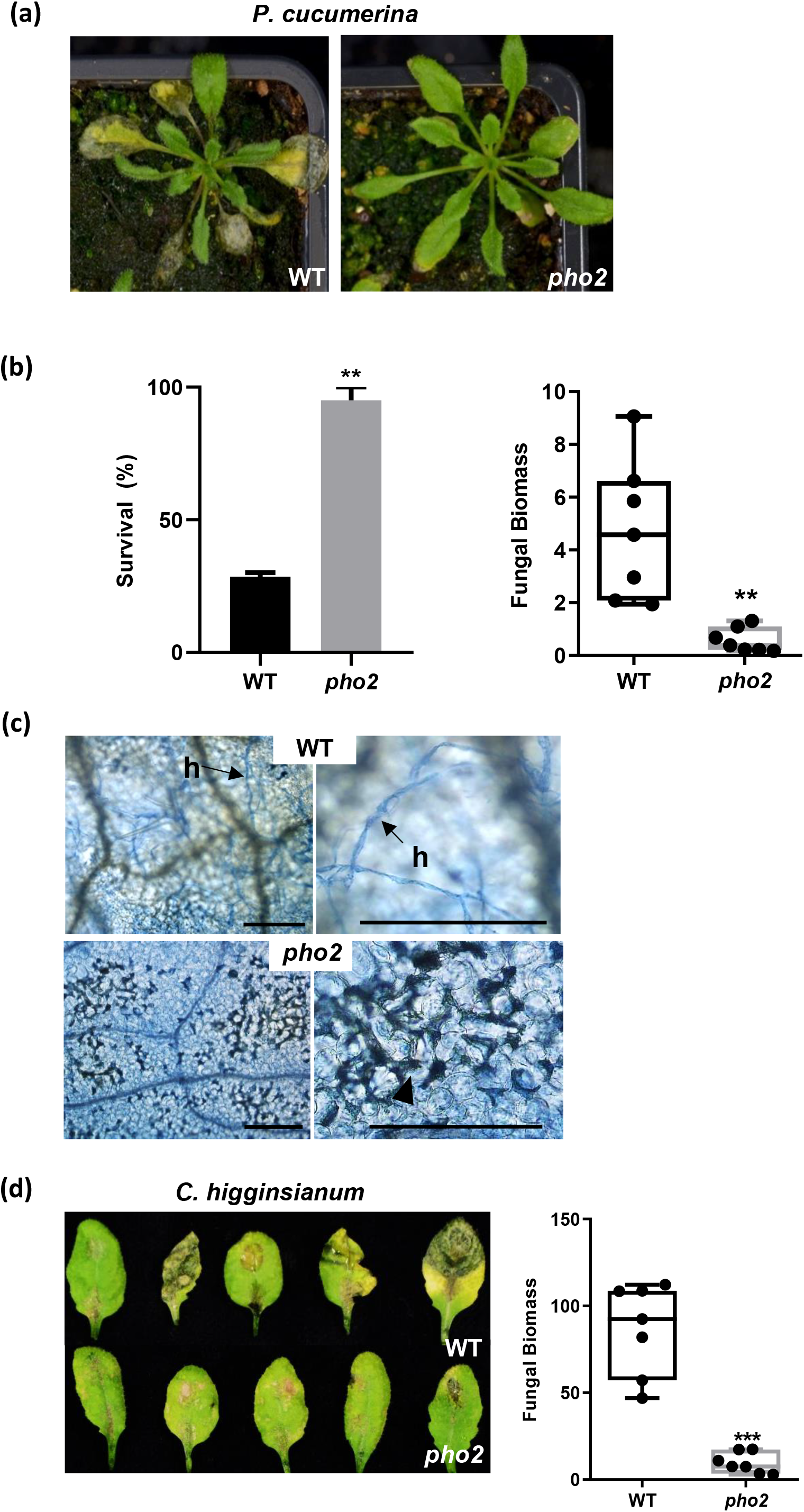
Resistance to infection by fungal pathogens in *pho2* mutant plants. Three week-old mutant plants were inoculated with fungal *P. cucumerina* spores. Three independent experiments were carried out with similar results with at least 12 plants per genotype. (a) Disease phenotype of wild type and *pho2* plants upon inoculation with *P. cucumerina* spores (5 x 10 spores/ml). Pictures were taken at 7 days post inoculation (dpi). (b) Survival ratio of WT and *pho2* plants at 7dpi (left panel). Quantification of *P cucumerina* DNA was carried out using specific primers of *P. cucumerina β-tubulin* at 7 dpi (right panel). Values are fungal DNA levels normalized against the Arabidopsis *UBIQUITIN21* gene (At5g25760). Data are mean± *SEM (n = 7) (t* test, \**p* ≤ 0.05, \*\**p* ≤ 0.01 and \*\*\**p* ≤ 0.001). (c) Trypan blue staining of *P. cucumerina*-infected leaves and visualization of cell death and fungal growth. h, hyphae. Arrows and arrowheads indicate fungal hyphae and cell death, respectively. Bars represent 200 μm. (d) Disease phenotype of wild type and *pho2* plants at 8 days after inoculation with *C. higginsianum* spores (4 x 10^6^ spores/ml).

### *MIR399* expression is up-regulated during fungal infection and treatment with fungal elicitors

To gather further support for the involvement of miR399 in Arabidopsis immunity, we investigated whether *MIR399* show changes in expression in response to infection with *P. cucumerina* in wild type plants. The accumulation of precursor transcripts and mature sequences for miR399 was examined at 48h and 72h post-inoculation (hpi) with *P. cucumerina* spores. An increase in the accumulation of both precursor and mature miR399 sequences was clearly observed in the response of wild type plants to pathogen infection (Figure 4a).

**Figure 4.**
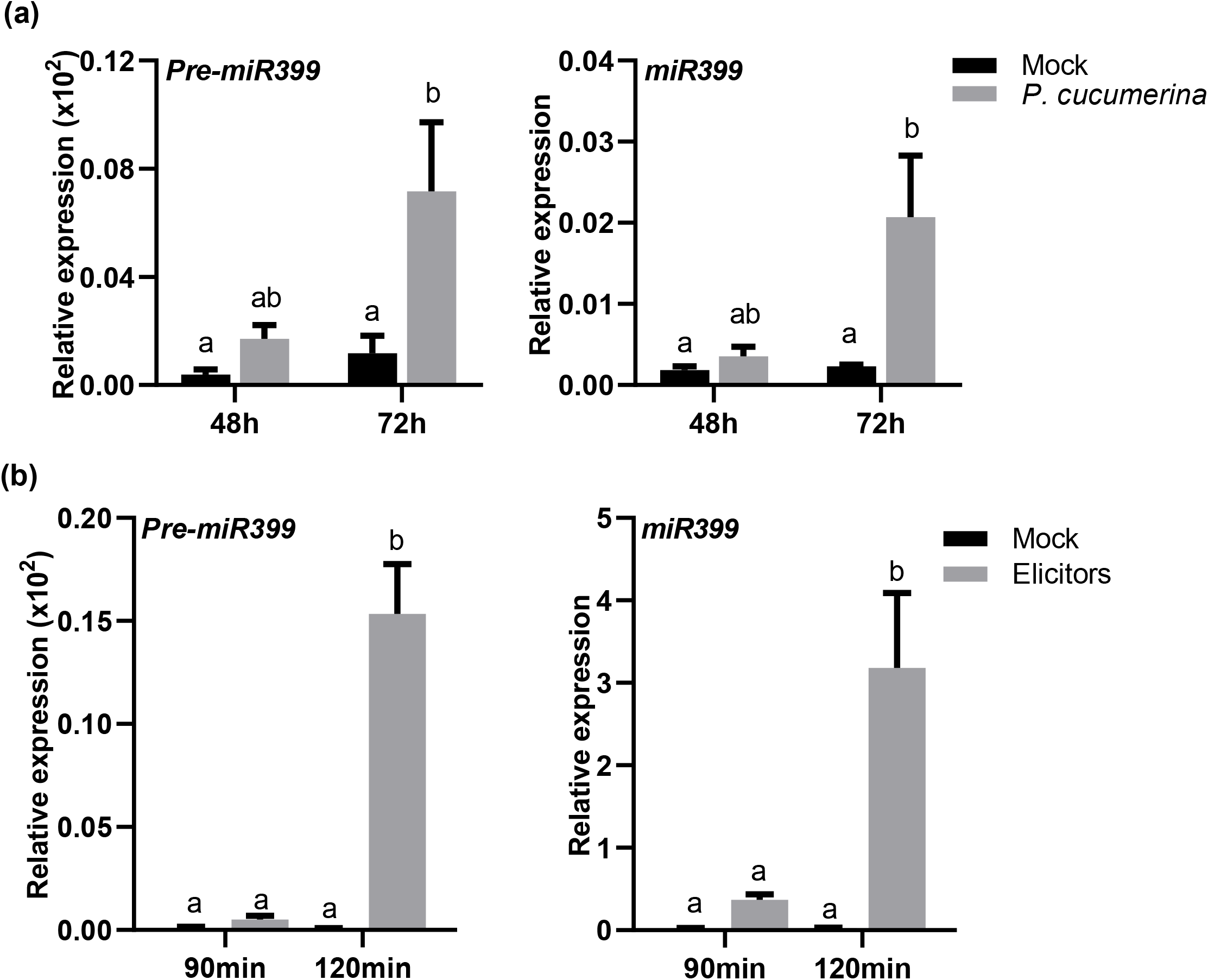
Accumulation of precursor (pre-miR399) and mature (miR399) transcripts upon inoculation with *P. cucumerina* spores (a) or treatment with *P. cucumerina* elicitors (b). Transcript accumulation was determined by RT-qPCR (pre-miR399) and stem-loop RT-qPCR (miR399) analysis at the indicated times after inoculation with fungal spores (a) or elicitor treatment (b). Four biological replicates and three technical replicates per time point were assayed. Statistically significant differences were determined by ANOVA, HSD Tukey’s test where different letters represent statistically significant differences.

In PTI, the activation of defense mechanisms relies on the detection of PAMPs, (commonly referred to as elicitors). Accordingly, we investigated whether miR399 accumulation is affected by treatment with a crude preparation of elicitors obtained by autoclaving and sonicating *P. cucumerina* mycelium. Similar to *P. cucumerina* infection, elicitor treatment induced both miR399 precursor and mature sequences (Figure 4b). The observation that pathogen infection and treatment with fungal elicitors is accompanied by up-regulation of *MIR399* expression suggest that miR399 might function in PTI in Arabidopsis.

### Stimulation of ROS production in Arabidopsis plants containing increased levels of Pi

One of the hallmarks of host-pathogen interactions is the overproduction of ROS as a plant defense mechanism, the so-called oxidative burst. ROS include various forms of reactive molecules, such as superoxide radicals (O_2_^.-^), hydroxyl radicals (OH^.^) or hydrogen peroxide (H_2_O_2_). Of them, H_2_O_2_, might act as both as signaling molecule for the activation of plant immune responses as well as an antimicrobial agent (Torres *et al.*, 2006). *RBOHD* (*Respiratory Burst Oxidase Homolog D*), a member of the Arabidopsis NADPH (Nicotinamide Adenine Dinucleotide Phosphate) oxidase gene family, has been shown to be responsible for ROS production after pathogen infection (Torres *et al.*, 2002; Kadota *et al.*, 2015). Furthermore, ROS may promote cell death and limitation of pathogen spread.

Knowing that miR399 OE and *pho2* plants exhibit a pattern of cell death upon pathogen infection, we examined ROS accumulation in miR399 OE and *pho2* plants, both in the presence and in the absence of pathogen infection. Histochemical detection of H_2_O_2_ was carried out using the fluorescent probe H_2_DCFDA (2’, 7’ dichlorofluorescein diacetate). H_2_DCFDA was previously shown to detect different forms of ROS, mainly H_2_O_2_ but also hydroxyl radical and superoxide anion (Fichman *et al*., 2019). Compared with wild type plants, a higher level of ROS accumulation could be observed in leaves of miR399 OE and *pho2* plants compared to wild type plants in the absence of pathogen infection (Figure 5a, upper panels). *P. cucumerina* infection further increased ROS levels in miR399 OE and *pho2* plants (Figure 5a, lower panels). Discrete regions accumulating ROS, most probably, correspond to infection sites. Similar results were observed by 3,3’-diaminobenzidine (DAB) staining of *P. cucumerina*-infected leaves (Figure S3). Moreover, H_2_DCFDA staining revealed a higher level of ROS accumulation in high Pi plants (e.g. miR399 OE and *pho2* plants) that have been treated with *P. cucumerina* elicitors compared with elicitor-treated wild type plants (Figure S4).

**Figure 5.**
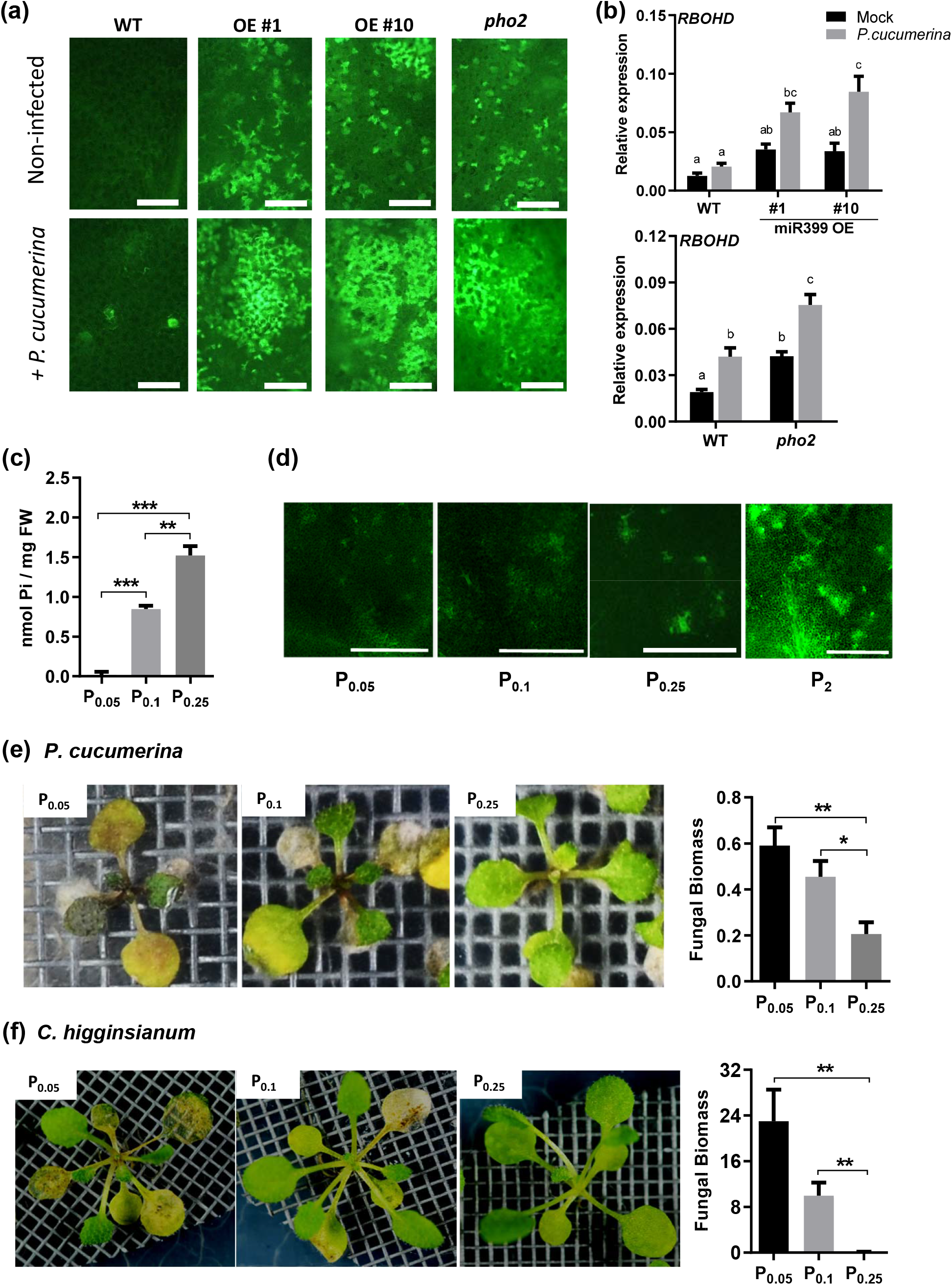
Enhanced accumulation of ROS and disease resistance in Arabidopsis plants that overaccumulate Pi. (a) *In situ* histochemical detection of ROS in leaves of miR399 OE and *pho2* plants. Plants were spray-inoculated with *P. cucumerina* spores (5 x 10 spores/ml), or mock-inoculated. Visualization of H_2_O_2_ accumulation was carried out using the fluorescent probe H_2_DCFDA at 2 days post inoculation (dpi). Bars represent 200 μm. (b) Expression of *RBOHD* in mock-inoculated and *P. cucumerina*-inoculated plants at 24hpi (black and grey bars, respectively). The expression values were normalized to the Arabidopsis *β-tubulin2* gene (At5g62690). Three biological replicates (with 3 plants per replicate) were examined. Different letters represent statistically significant differences (ANOVA, HSD Tukey’s test; P < 0.05). (c) Free Pi content in plants that have been grown under different conditions of Pi supply. Plants were grown on agar plates for 1 week, transferred to fresh agar plates with medium containing different concentrations of Pi (0.05mM, 0.lmM, or 0.25mM). Plants were allowed to continue growth for one more week and then inoculated with fungal spores. Pi content was determined at the time of inoculation with fungal spores. (d) ROS accumulation in wild type Arabidopsis (Col-0) plants that have been grown under different Pi supply conditions that is 0.05mM, 0.1mM, 0.25mM, or 2 mM Pi (P_0.05_, P_0.1_, P_0.25_, and P_2_, respectively). ROS was detected using H_2_DCFDA. Representative images are shown. Bars correspond to 1 mm. (e) Resistance to infection by *P. cucumerina* in Pi-treated Arabidopsis plants. Appearance of plants at 7 days post-inoculation (dpi) with *P. cucumerina* spores (4 x 10 spores/ml; left panel). Representative results from one of three independent infection experiments that gave similar results are shown. Right panel, fungal biomass determined at 3 dpi by quantitative PCR analysis using specific primers of *P. cucumerina β-tubulin* and normalized to Arabidopsis *UBIQUITIN21* gene (At5g25760). (f) Resistance to infection by *C. higginsianum* in Pi-treated Arabidopsis plants. Disease symptoms of Arabidopsis plants at 12 days post-inoculation with *C. higginsianum* spores (5 x 10^5^ spores/ml). Right panel, fungal biomass determined at 7 dpi by quantitative PCR analysis using specific primers of *C. higginsianum ITS2 (Internally transcribed spacer 2).* Means of three biological replicates, each one from a pool of at least three plants are shown in e and f (right panels; *t* test, \**p* ≤ 0.05, \*\**p* ≤ 0.01 and \*\*\**p* ≤ 0.001).

Additionally, we examined ROS generated in miR399 OE, *pho2* plants and wild type plants in response to treatment with *P. cucumerina* elicitors using the luminol method. This study confirmed a higher production of ROS after elicitor treatment in leaves of miR399 OE and *pho2* plants compared to that in wild type plants (Figure S5a). Collectively, these findings support ROS accumulation in leaves of miR399 OE and *pho2* plants in the absence of pathogen infection. During infection with *P. cucumerina,* or treatment with elicitors obtained from this fungus, miR399 OE and *pho2* plants produced higher levels of ROS than wild type plants.

In concordance with results obtained by histochemical detection of ROS and measurement of ROS production using the luminol assay, the miR399 OE and *pho2* mutant plants exhibited up regulation of *RBOHD* in the absence of pathogen infection compared with wild type plants (Figure 5b, black bars). *P. cucumerina* infection further induced *RBOHD* expression in all the genotypes (WT, miR399 OE and *pho2* plants), but its expression reached higher levels in miR399 OE and *pho2* plants than in wild type plants (Figure 5b, grey bars). Then, ROS accumulation in miR399 OE and *pho2* plants can be explained by an increased *RBOHD* expression. However, other factors causing an increase in ROS accumulation cannot be discarded.

The observation that both miR399 OE and *pho2* plants accumulated Pi and ROS, these plants also exhibiting enhanced disease resistance (Figure 1 and 2), prompted us to investigate whether ROS production and disease resistance is affected by Pi supply in Arabidopsis. To address this question, wild type plants were grown *in vitro* on media at different Pi concentrations (0.05 mM, 0.1 mM, 0.25 mM Pi, hereinafter referred as P_0.05_, P_0.1_ and P_0.25_ plants). As expected, measurement of Pi content confirmed that increasing Pi supply to wild type plants results in higher leaf Pi content (Figure 5c). Most importantly, increasing Pi supply was accompanied by an increase in ROS accumulation as revealed by H_2_DCFDA staining of leaves from Pi-treated plants (Figure 5d). Similarly, luminol assays demonstrated that treatment with Pi fosters ROS production in wild type plants, with the greater ROS production correlating well with the higher Pi concentration (Figure S5b). Most importantly, upon pathogen inoculation, high-Pi plants consistently displayed enhanced resistance to infection by either *P. cucumerina* or *C. higginsianum* (Figure 5e, f, respectively).

### Pi-induced resistance to pathogen infection in Arabidopsis occurs through modulation of SA- and JA-dependent defense pathways

As previously mentioned, SA and JA play a critical role in the transcriptional reprograming of Arabidopsis plants in response to pathogen infection (Pieterse *et al.*, 2012). A pathogen-induced accumulation of ROS is also required for the induction of SA-dependent defenses, indicating that ROS and SA are intertwined in a complex regulatory network (Wang *et al.*, 2014; Herrera-Vásquez *et al.*, 2015). To get deeper insights into the mechanisms underlying disease resistance in miR399 OE and *pho2*, we investigated the expression of defense genes linked to the SA and the JA pathways in these plants. The marker genes of the SA-mediated defense response here examined were: *PR1* (*Pathogenesis-Related 1*), *NPR1* (*Non-expressor of Pathogenesis-Related genes 1*), and *PAD4* (*Phytoalexin Deficient4*) (Jirage *et al.*, 1999). We found that, in the absence of pathogen infection, *PR1*, *NPR1* and *PAD4* expression was up-regulated in both miR399 OE and *pho2* plants compared with wild type plants (Figure 6a, black bars). *PR1*, *NPR1* and *PAD4* expression further increased in response to *P. cucumerina* infection in all the genotypes (Figure 6a, grey bars).

**Figure 6.**
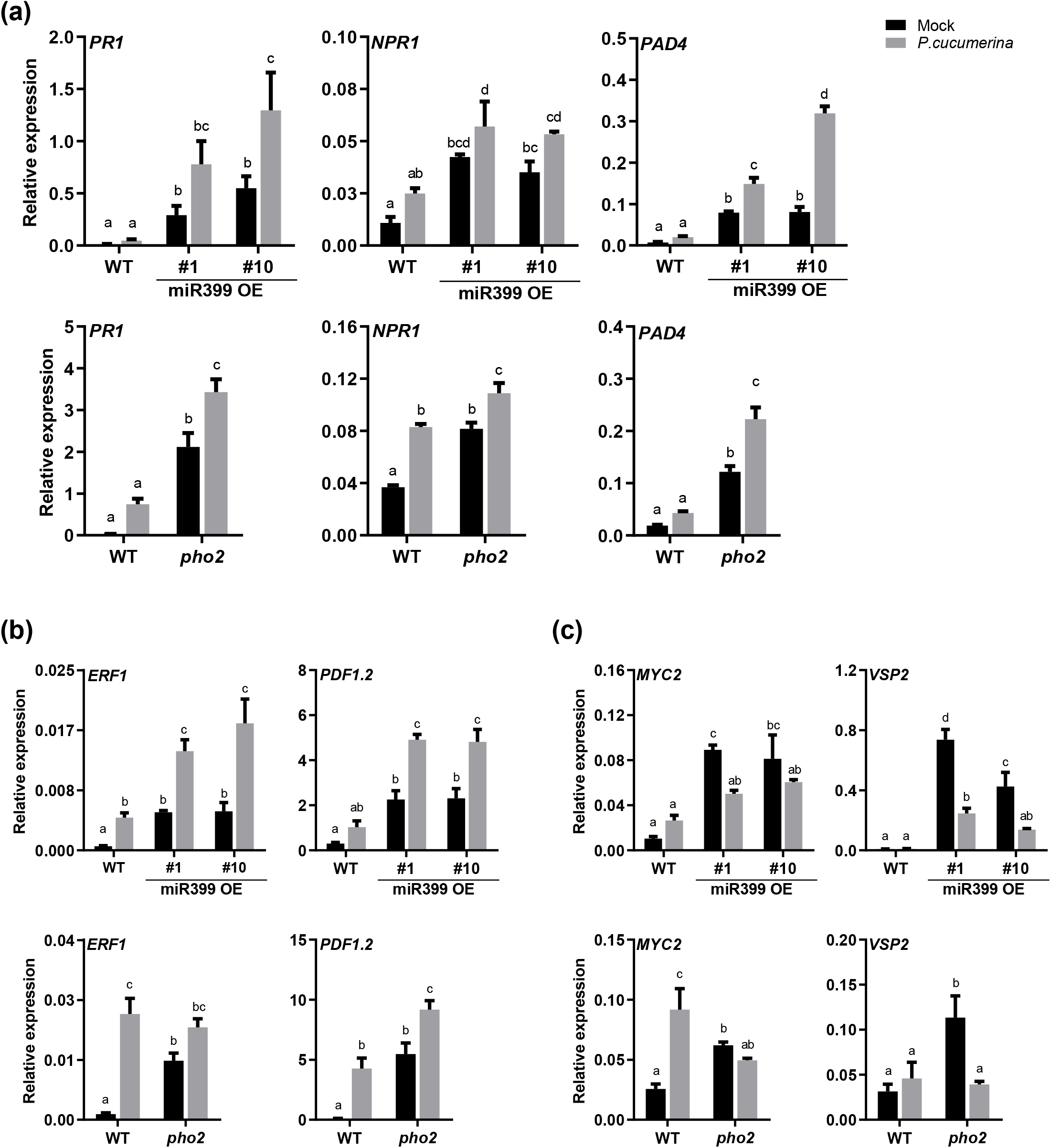
Expression of defense genes in miR399 OE and *pho2* mutant plants. Transcript levels were determined by RT-qPCR in mock-inoculated or *P. cucumerina*-inoculated plants at 24 hours post-inoculation (black and grey bars, respectively). The expression values were normalized to the Arabidopsis *β-tubulin2* gene (At5g62690). (a) Expression of genes involved in the SA pathway (*PR1, NPR1, PAD4).* (b) Expression of genes in the ERF branch of the JA pathway (*ERF1, PDF 1.2*). (c) Expression of genes in the MYC branch of the JA pathway (*MYC2, VSP2).* Three independent experiments (with 12 plants per genotype) were examined, with similar results. Bars represent mean ± SEM. Different letters represent statistically significant differences (ANOVA, HSD Tukey’s test; P < 0.05).

Regarding the JA pathway, two branches are documented in Arabidopsis: the MYC2 branch that is regulated by AtMYC2 (a basic helix-loop-helix-leucine zipper transcription factor), and the ERF branch, which is regulated by AtERF1 (a member of the APETALA/ERF transcription factor family) (Lorenzo *et al.*, 2004). PDF1.2 (*Plant Defensin 1.2*) is commonly used as marker of the ERF branch, whereas the MYC branch is marked by the induction of *VSP2 (Vegetative Storage Protein 2*) (Lorenzo *et al.*, 2004; Pieterse *et al.*, 2012; Wasternack and Hause, 2013; Zhang *et al.*, 2017). The ERF branch and the MYC branch of the JA signaling pathway have been reported to repress each other (Lorenzo *et al.*, 2004; Wasternack and Hause, 2013; Aerts *et al.*, 2021).

Compared with wild type plants, the miR399 OE and *pho2* plants showed higher expression of transcription factor and defense marker genes associated to the two branches of the JA signaling pathway in the absence of pathogen infection, namely the ERF1/PDF1.2 and MYC2/VSP2 branches (Figure 6b, c, black bars). Pathogen infection further induced *ERF1*/*PDF1* expression in all the genotypes (wild type, miR399 OE and *pho2* plants) (Figure 6b, grey bars). To note, while *MYC2*/*VSP2* expression was induced by pathogen infection in wild type plants, their expression was repressed both in *P. cucumerina*-infected miR399OE and *P. cucumerina*-infected *pho2* plants (compared with the corresponding non-infected plants) (Figure 6c, grey bars). Together, these findings support that resistance to pathogen infection in plants accumulating Pi (i.e. miR399 OE, *pho2* plants) is associated with a higher expression of SA- and JA-regulated genes under non-infection conditions. A pathogen-induced superactivation of genes involved in the SA and the ERF1 branch of the JA pathway occurs in these plants, while pathogen infection represses the MYC2 branch of the JA signaling pathway.

Further, we measured levels of the phytohormones SA and JA in miR399 OE and *pho2* plants. Compared with wild type plants, miR399 OE and *pho2* plants accumulated significantly higher levels of SA under normal growth conditions (e.g. in the absence of pathogen infection) (Figure 7a, upper left panel, black bars). This observation is consistent with the expression pattern of SA-responsive defense genes found in miR399 OE and *pho2* plants (see Figure 6). Upon pathogen infection, however, there were no significant differences in SA levels among wild type, miR399 OE and *pho2* plants (Figure 7a, upper left panel, grey bars). We also noticed that the SA glucoside SAG (the storage from SA) accumulated at substantially higher levels in the fungal-infected miR399OE and *pho2* plants compared with the fungal-infected wild type plants (Figure 7a, upper right panel). This observations points to a strict control in the SA level in miR399OE and *pho2* plants

**Figure 7.**
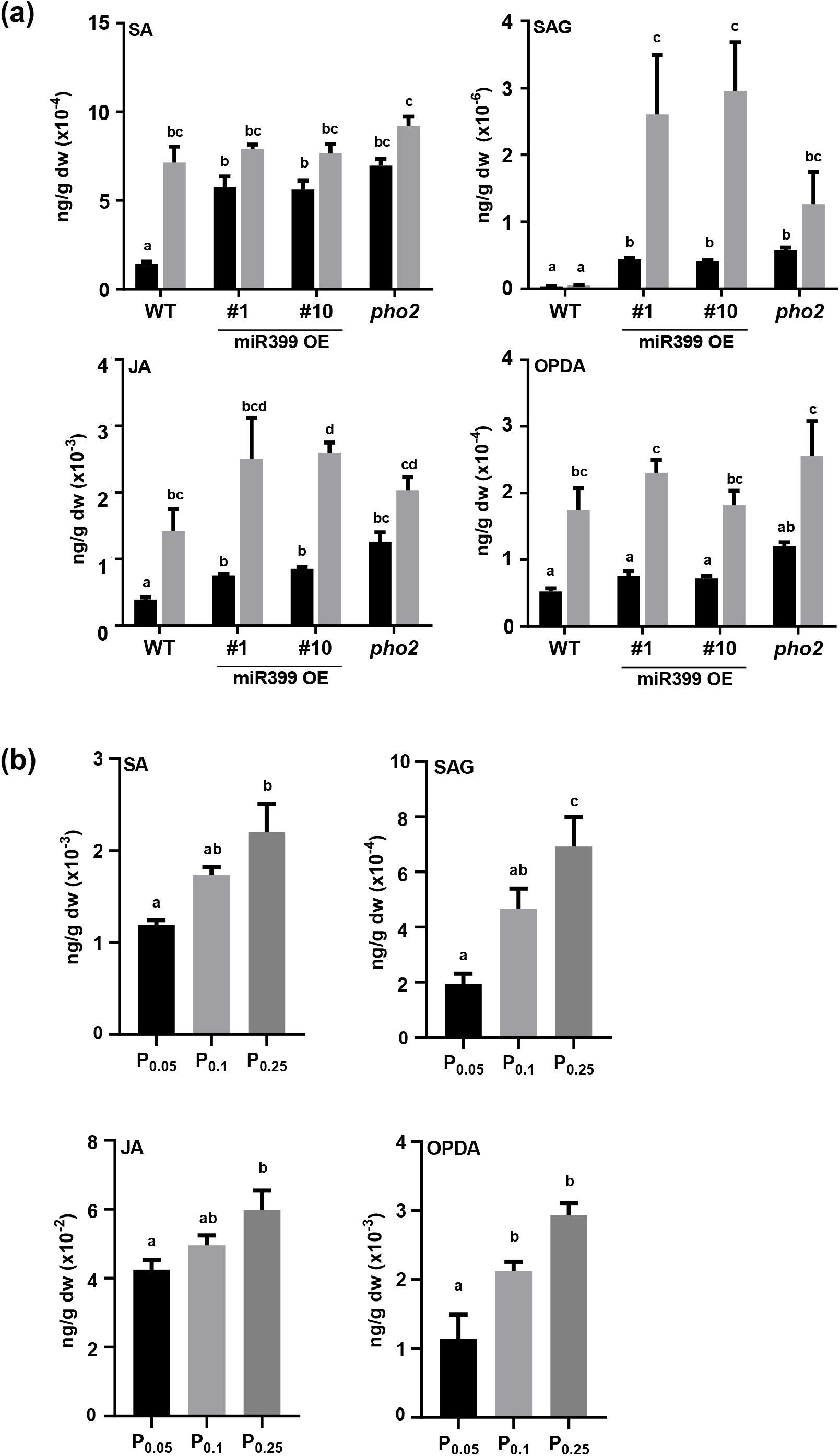
Levels of SA, SAG, JA and OPDA in leaves of Arabidopsis plants that overaccumulate Pi (miR399 OE, *pho2,* and wild type plants that have been grown under different Pi supply conditions). (a) Hormone levels were measured in mock-inoculated and *P. cucumerina-inoculated* miR399 and *pho2* plants at 48hpi. (b) Hormone levels in wild type plants that have been grown at the indicated Pi supply conditions (0.05, 0.1, and 0.25 mM Pi). Plants were grown as indicated in Figure 5. Three independent experiments (with 12 plants per genotype) were examined, with similar results. Bars represent mean ± SEM. Different letters represent statistically significant differences (ANOVA, HSD Tukey’s test; P < 0.05).

In the absence of pathogen infection, miR399 overexpression and loss-of-function of *PHO2* plants accumulated more JA than wild type plants, which was more evident in *pho2* plants (Figure 7a, lower left panel, black bars). JA content further increased during pathogen infection in all the genotypes (Figure 7a, lower left panel, grey bars). Regarding OPDA (12-oxophytodienoic acid), the JA biosynthetic precursor, its level in miR399 OE and *pho2* plants did not differ significantly from that in wild type plants (Figure 7a).

Additionally, we investigated whether Pi treatment modulates SA and JA signaling pathways. Increasing Pi supply results in a gradual increase in SA and JA level (as well as SAG and OPDA level) (Figure 7b). These results are in concordance with upregulation of SA- and JA-regulated defense genes (*PR1, NPR1* and *PDF1.2*) in response to Pi treatment (Figure S6).

Collectively, these results support that, in the absence of pathogen infection, miR399 overexpression, loss-of-function of *PHO2*, and Pi treatment results in a higher level of SA and JA, and subsequent up-regulation of SA- and JA-dependent defense gene expression. A Pi-mediated modulation of SA and JA signaling pathways might well explain the phenotype of disease resistance that is observed in Arabidopsis plants overaccumulating Pi, namely miR399 OE, *pho2*, and wild type plants grown under high Pi supply.

## DISCUSSION

Nutrients play crucial roles in normal plant growth and development, and nutrient imbalance might also have a substantial impact on the predisposition of plants to resist pathogen attack. Depending on the identity of the interacting partners, nutritional imbalances caused by either nutrient excess or deficiency may determine the outcome of the interaction, resistance or susceptibility (Veresoglou *et al.*, 2013). On the other hand, although miRNA-mediated regulation of gene expression in processes involved in either nutrient stress or immune responses is well documented, less effort has been made to investigate miRNA function in crosstalk between pathogen-induced and nutrient pathways, in particular Pi.

In this study, we provide evidence that increasing Pi content in Arabidopsis results in enhanced disease resistance. Several pieces of evidence support this conclusion. On the one hand, we show that Pi accumulation caused by miR399 overexpression, loss-of-function of *PHO2*, or treatment with Pi, confers resistance to infection by fungal pathogens with necrotrophic (*P. cucumerina*) and hemibiotrophic (*C. higginsianum*) lifestyle. On the other hand, resistance in Arabidopsis plants overaccumulating Pi correlated well with up-regulation of defense-related genes under normal growth conditions (i.e. in the absence of pathogen infection). Altogether, these findings support that miR399 functions as a regulator of Arabidopsis immunity, and reinforce the notion that miR399 plays a dual role in plants by controlling Pi homeostasis and immune responses. Further supporting a role for miR399 in Arabidopsis immunity, we show that *P. cucumerina* infection is accompanied by an increase in miR399 accumulation. Not only pathogen infection, but also treatment with fungal elicitors results in higher levels of miR399 indicating that MIR399 is PAMP-responsive and might function in PTI.

Another finding of this study is that, under non-infection conditions, overaccumulation of Pi in Arabidopsis leaves is accompanied by production of ROS, its level increasing significantly during pathogen infection. Hence, ROS accumulation in miR399OE and *pho2* plants might be responsible for the pattern of cell death observed in these plants during pathogen infection, a response that is reminiscent of the pathogen-induced HR. In contrast, the fungal-infected wild type plants did not exhibit cell death and showed extensive fungal growth. Additionally, elevated levels of ROS might contribute to the activation of immune responses in high Pi plants leading to a phenotype of disease resistance. From this point of view, it might be interesting to determine whether Pi-induced ROS accumulation can be generalized to other plant species.

At the time of inoculation with fungal spores (three-week-old plants), miR399OE and *pho2* plants accumulated 3-4 times more Pi than WT plants. By this developmental stage, plant growth is not compromised in plants accumulating this level of Pi. It is tempting to hypothesize that this increase in Pi content and/or ROS level is perceived by the host plant as a stressful situation, and that the plant responds to these signals with the induction of defense gene expression. An increase in Pi content might eventually increase intracellular and extracellular Pi levels for an increase in ATP level. Here, it should be mentioned that ATP has been shown to function as a DAMP signal after release into the extracellular space upon cellular damage, and that extracellular ATP enhances plant defense against pathogens through the activation of JA (Tanaka *et al.*, 2014; Tripathi *et al.*, 2018). Further studies are needed to establish whether ATP levels are altered in high-Pi plants.

Gene expression analysis revealed regulation of the SA- and JA-defense signaling pathways in Arabidopsis plants overaccumulating Pi in leaves. Compared to wild type plants, all the high Pi plants here examined (miR399 OE, *pho2,* and plants grown under high Pi supply conditions) showed activation of SA-regulated (*PR1, NPR1, PAD4*) and JA-regulated (*ERF1* and *PDF1.2; MY2* and *VSP2*) genes under non-infection conditions. SA and JA were reported to play a positive role in the regulation of resistance to *P. cucumerina* infection in Arabidopsis (Berrocal-Lobo *et al.*, 2002; Sánchez-Vallet *et al.*, 2012). Previous studies also revealed that *PDF1.2* induction was associated to resistance to infection by necrotrophic fungi, including *P. cucumerina*, in Arabidopsis (Thomma *et al.*, 1998; Berrocal-Lobo *et al.*, 2002). Consistent with upregulation of genes involved in the SA and JA signaling pathways, SA and JA accumulated in non-infected high Pi plants. In other studies, a feed-forward loop between SA and ROS production (e.g. H_2_O_2_) has been reported in which ROS are involved both upstream and downstream of SA in the plant defense response to pathogen infection (Herrera-Vásquez *et al.*, 2015). We hypothesize that Pi content and, possibly also ROS accumulation, might influence defense hormone signaling.

Interestingly, a different regulation in the two branches of the JA pathway was observed in high-Pi plants during pathogen infection. Whereas *ERF1* and *PDF1.2* expression is further increased during infection in high Pi plants (infected vs noninfected plants, each genotype), *MYC2* and *VSP2* expression diminished in these plants (infected vs non-infected plants, each genotype). We envisage that this differential regulation might be due to still unknown factors that cooperate in an antagonistic manner in the regulation of the ERF1 and MYC2 branches of the JA pathway during infection with the fungal pathogen *P. cucumerina*. In line with this, a negative correlation between the ERF1 and MYC2 branches has been previously described in the Arabidopsis response to different attackers such as pathogens and phytophagous insects (Lorenzo *et al.*, 2004; Pieterse *et al.*, 2012; Wasternack and Hause, 2013; Zhang *et al.*, 2017). Here, necrotrophic pathogens preferentially activate the ERF branch, while the MYC2 branch is activated by insect herbivory and wounding. A specialization in the host plant for modulation of each branch of the JA signaling pathway might then exists. It will be of interest to explore whether Pi accumulation has an effect on plant/insect interactions in Arabidopsis.

Clearly, cross-talk between defense-related hormone pathways provides the plant with a powerful regulatory potential to control defense responses (Zheng *et al.*, 2012; Aerts *et al.*, 2021). However, the type of induced response that is effective for disease resistance appears to vary depending on the host plant and pathogen identity. Although there are exceptions, pathogens with a biotrophic or hemibiotrophic lifestyle (such as *Pseudomonas syringae*) are generally more sensitive to SA-dependent responses, whereas necrotrophic pathogens are commonly deterred by JA/ET-dependent defenses (Glazebrook, 2005). The observation that high Pi plants (e.g. miR399 OE, *pho2,* and plants grown under high Pi supply conditions) show enhanced resistance to necrotrophic (*P. cucumerina*) and hemibiotrophic (*C. higginsianum*) fungal pathogens makes it unlikely that the pathogen lifestyle determines disease resistance in high Pi plants.

Based on the results obtained in this study, a model is proposed to describe possible mechanisms underlying disease resistance in high Pi Arabidopsis plants (Figure 8). According to our model, miR399 overexpression and loss-of-function of *PHO2,* as well as growing plants under high Pi supply, increases Pi accumulation, ROS production, as well as SA and JA accumulation. A higher expression of SA- and JA-dependent defense responses in Arabidopsis plants accumulating Pi would allow the plant to mount a successful defense response upon encountering a pathogen.

**Figure 8.**
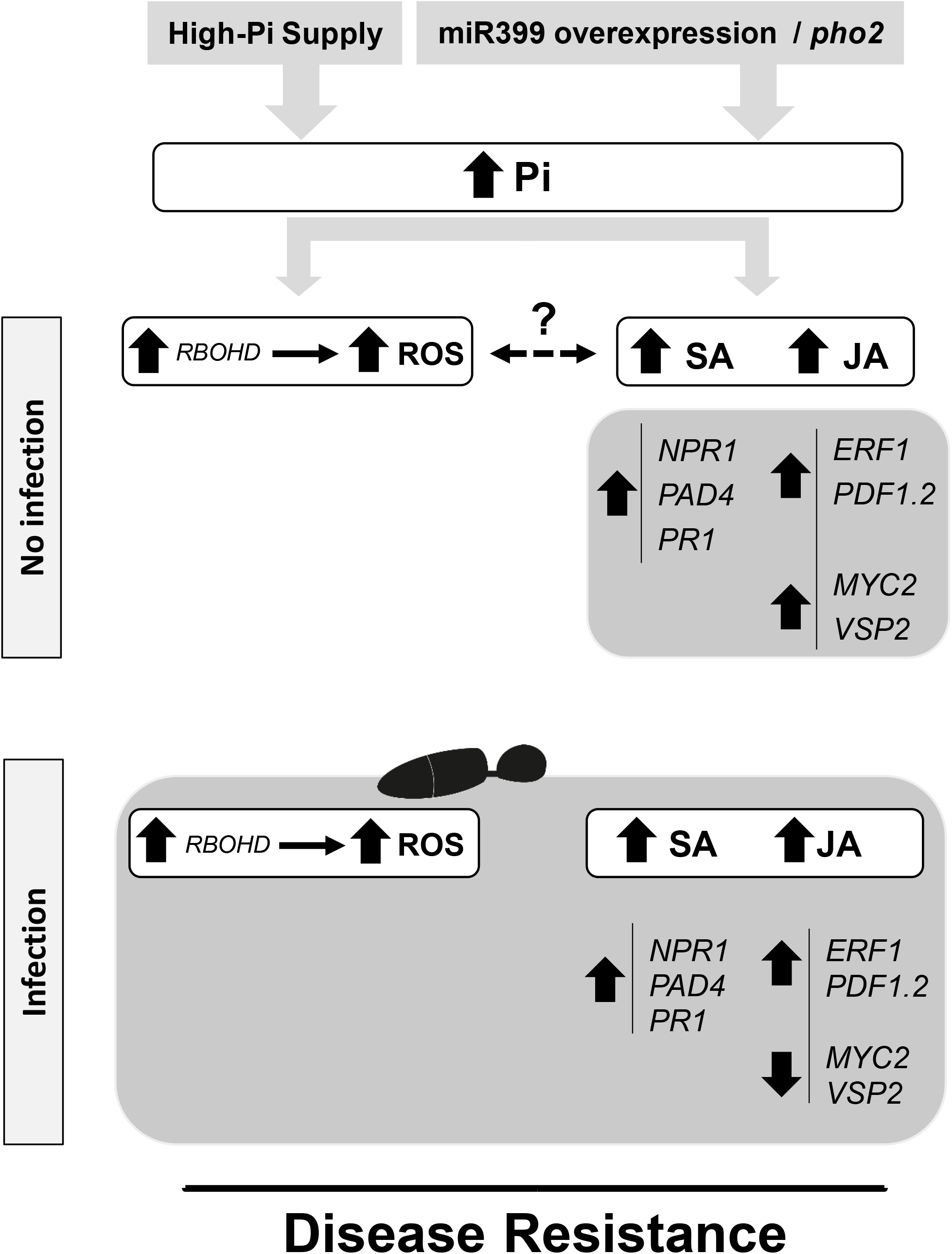
Proposed model to explain how Pi content and defense responses are integrated for modulation of resistance to infection by fungal pathogens in Arabidopsis. Treatment with high Pi, *MIR399* overexpression and loss-of-function of *pho2* would trigger Pi accumulation which, in turn, would increase ROS and hormone (SA and JA) levels in Arabidopsis leaves for the induction of genes involved in the SA and JAsignaling pathway. Upon pathogen infection, the expression of genes involved SA and *ERF1/PDF1.2* branch of the JA pathway would be further induced, while genes in the *MYC2/VSP2* branch of the JA pathway are repressed. The interplay between Pi, ROS and hormone would allow the plant to mount an effective immune response. Although crosstalk between ROS and hormonal pathways have been described (Herrera-Vásquez *et al.*, 2015; Xia *et al.*, 2015), the exact mechanisms by which Pi content modulates ROS production and hormone content, and how these signaling pathways cross-talk to each other, in the coordinated regulation of Pi homeostasis and immune responses deserves further investigation.

The existence of links between components of the phosphate starvation response and disease resistance has been previously described (Chan *et al.*, 2021). For instance, enhanced resistance to infection by the bacterial pathogen *P. syringae* DC3000 and the oomycete pathogen *Hyaloperonospora arabidopsidis* was reported in *phr1* mutant plants, *PHR1* being the master transcriptional regulator of the Arabidopsis Phosphate Starvation Response (PSR) (Castrillo *et al.*, 2017). This study also led the authors to propose that *PHR1* might fine-tune JA responses under Pi starvation in specific biological contexts, rather than being a regulator of the JA signaling pathway (Castrillo *et al.*, 2017). In other studies, transgenic expression of a phytoplasma effector (SAP11) in Arabidopsis was found to trigger Pi starvation responses that are mainly dependent of *PHR1* (Lu *et al.*, 2014). The SAP11 transgenic plants overaccumulated Pi in leaves and were more susceptible to *P.syringae* pv.tomato DC3000 infection (Lu *et al.*, 2014). The PSR system also appears to control root colonization by the endophytic fungus *Colletotrichum tofieldiae* in Arabidopsis (Hiruma *et al.*, 2016; Frerigmann *et al.*, 2021). In citrus plants, the Pi content was associated to symptomology in the Huanglongbing (HBL) disease, where Pi deficiency favors development of HLB symptoms (Zhao *et al.*, 2013). In plant-insect interactions, Pi deficiency was found to induce the JA signaling pathway to enhance resistance to insect herbivory in a process that is partially under the control of *PHR1* (Khan *et al.*, 2016). Collectively, results here presented together with those found in the literature in other pathosystems support the existence of connections between Pi and immune signaling. These mechanisms should operate in a coordinated manner to properly balance nutrient responses and plant immunity.

Results from our study also raise intriguing questions about the impact of Pi content in disease resistance in different pathosystems. Thus, in this study we show that an increase in Pi content positively regulates Arabidopsis immune responses which, in turn, enhances resistance to infection by fungal pathogens. Contrarily, in rice plants, a higher Pi content caused by miR399 overexpression or high Pi fertilization was found to negatively regulate defense gene expression, thus, increasing susceptibility to infection by the blast fungus *Magnaporthe oryzae* (Campos-Soriano *et al.*, 2020). In other studies, Pi deficiency was found to enhance resistance to *Verticillium dahliae* in cotton (Luo *et al.*, 2021) and insect herbivory in Arabidopsis (Khan *et al.*, 2016). Together, our results and those previously reported support that Pi content might positively or negatively regulate disease resistance depending on the interacting partners. It is tempting to speculate that different plants might have evolved diverse mechanisms to adapt to Pi alterations which, in turn, would determine a different effect of Pi in the regulation of immune responses. It is likely that integration of Pi stress (either deficiency or excess) and immune responses might vary depending on the host plant and the type of pathogen, the outcome of the interaction being also dependent on the role of the defense hormones SA and JA in that particular interaction. Alternatively, Pi might affect growth and/or pathogenicity of a fungal pathogen, either by creating a less favorable environment for pathogen growth, or by reducing the production of pathogen virulence factors. Clearly, these aspects deserve further investigation.

The fact that miR399 plays a positive role in regulating immune responses in Arabidopsis (present work), but a negative role in the regulation of immune responses in rice (Campos-Soriano *et al.*, 2020) is not unusual. In the literature, there are examples of miRNAs with an opposite role in disease resistance depending on the plant/pathogen interaction. For instance, miR398 overexpression in Arabidopsis compromises resistance against the bacterial pathogen *P. syringae*, while its overexpression in rice enhances resistance to the blast fungus *M. oryzae* (Li *et al.*, 2010; Li *et al.*, 2019). These observations indicate that a particular miRNA might function either as a positive or as a negative regulator of immune responses depending on the host plant and the type of pathogen.

Finally, it is worth mentioning that one of the major challenges plants face is defending against pathogen infection under continuous changes in nutrient availability, particularly Pi availability. Results here presented set the basis for further research to elucidate the exact mechanisms by which Pi signaling and pathogen-induced signaling interact with each other in plants. Further studies, however, should be conducted on a case-by-case basis in different plant/pathogen interactions. A better understanding of these mechanisms will allow the development of novel strategies to improve disease resistance in plants.

## EXPERIMENTAL PROCEDURES

### Plant material and infection assays

*Arabidopsis thaliana* (ecotype Columbia-0) plants were grown in a mixture of soil:perlite:vermiculite (2:1:1) and modified Hoagland half strength medium, under neutral photoperiod (12h light / 12h dark), 60% of humidity and a temperature of 22°C ± 2°C. The fungus *Plectosphaerella cucumerina* was grown on PDA (Potato Dextrose Agar) plates with chloramphenicol (34 μg/ml). *Colletotrichum higginsianum* was grown on Oatmeal agar plates in darkness. Fungal spores were collected by adding sterile water to the surface of the mycelium, and adjusted to the desired final concentration using a Bürker counting chamber.

For infection experiments in soil-grown plants, three-week-old plants were spray-inoculated with a spore suspension of *P. cucumerina* (5 x 10^5^ spores/ml), or mock-inoculated. *P. cucumerina*-inoculated and mock-inoculated plants were maintained under high humidity and plant survival was assessed at 7 dpi. For infection with *C. higginsianum,* the fungal spores were locally-inoculated (4 x 10^6^ spores/ml; 10 μl/leaf and 5 leaves/plant). Lesion area of *C. higginsianum*-infected leaves was measured with software ImageJ (National Institute of Health, Bethesda, MD, USA; https://imagej.nih.gov/ij/) at 7 dpi. Three independent experiments were performed with at least 12 plants per genotype in each experiment. For *in vitro* experiments, two-week old Arabidopsis plants were spray-inoculated with *P. cucumerina* (4 x 10^6^ spores/ml) or locally-inoculated with *C. higginsianum* (5 x 10^5^ spores/ml). Fungal biomass was quantified by real-time PCR using specific primers for the corresponding fungus and the Arabidopsis *UBIQUITIN21* (At5g25760) gene as the internal control (Soto-Suárez *et al.*, 2017). PCR primers are listed in Table S1. Statistically significant differences were determined by t-test.

Elicitor treatments were performed by spraying three-week old plants with an elicitor extract obtained from the fungus *P. cucumerina* (300 μg/ml) as previously described (Casacuberta *et al.*, 1992). Three independent experiments were performed with at least 12 plants per genotype in each experiment.

For Pi treatment experiments, plants were grown *in vitro* on meshes placed on agar plates with modified Hoagland half strength medium containing 0.25 mM KH_2_PO_4_ for one week. Seedlings were then transferred to fresh agar-medium at the desired concentration of Pi (0.05, 0.1, 0.25, or 2 mM Pi). The plants were allowed to continue growing for one more week under each Pi regime. The *in vitro*-grown plants were then inoculated with a spore suspension of *P. cucumerina* or *C. higginsianum* as above.

The *pho2* mutant, previously named *UBC24* (UBIQUITIN-CONJUGATING ENZYME 24 (At2g33770; Columbia background) was obtained from the Arabidopsis Biological Resource Center (ABRC, ref. CS8508). A point mutation in the sixth exon (from G_2539_ to A, relative to the translational start codon) causes an early termination at the beginning of the UBC domain, thus, resulting in the loss of the ubiquitin-conjugating activity of PHO2 in the *pho2* mutant (Aung *et al.*, 2006).

### Plant Tissue staining

For trypan blue staining, leaves were fixed by vacuum infiltration for 1h in ethanol:formaldehyde:acetic acid (80:3.5:5 v/v), stained with lactophenol blue solution for one hour, and then washed with chloral hydrate for 15 minutes. Leaves were placed on glass slides with glycerol and observed using a Leica DM6 microscope under bright field.

For H_2_DCFDA staining, the Arabidopsis leaves were placed on a solution of H_2_DCFDA (at a concentration of 10μM), vacuum infiltrated during 5 minutes, and then maintained in darkness for 10 minutes. Two washes with distillated water were performed. Photographs were taken on a Leica DM6 microscope to visualize green fluorescence. Diaminobenzidine tetrahydrochloride (DAB) staining was also used to examine H_2_O_2_ levels. For this, Arabidopsis plants were immersed in a DAB solution (1 mg/ml) for 30 minutes with vacuum treatment, maintained during 4 hours in darkness and agitation, washed with 95% ethanol for 30 minutes at 75°C, and observed using a Leica DM6 microscope under bright-field illumination.

ROS production in response to treatment with *P. cucumerina* elicitors (300 μg/ml) was measured using the luminol method described by Smith and Heese (2014). Measurements were carried out on a CentroXS3 LB 960 Microplate (Berthold Technologies, Germany).

### Generation of transgenic Arabidopsis plants

For *MIR399f* overexpression, the miR399 precursor sequence was cloned under the control of the *Cauliflower mosaic virus (CMV) 35S* promoter in the PMDC32 plasmid (Chiou *et al.*, 2006). The plant expression vector was transferred to the *Agrobacterium tumefaciens* strain GV301. Arabidopsis (Col-0) plants were transformed using the floral dip method. Homozygosis was obtained by antibiotic selection. Segregation analysis confirmed transgene inheritance in successive generations of transgenic lines.

### Measurements of Pi content and chlorophyll content

The Pi content of Arabidopsis plants was determined as previously described (Versaw and Harrison, 2002). Chlorophylls were extracted and quantified spectrophotometrically (SpectraMax M3, Molecular Devices, CA, USA) as previously described (Lichtenthaler and Buschmann, 2001).

### Gene expression analyses

Total RNA was extracted using TRIzol reagent (Invitrogen). First-strand cDNA was synthesized from DNAse-treated total RNA (1 μg) with reverse transcriptase and oligo-dT (High Capacity cDNA reverse transcription kit, Applied Biosystems). RT-qPCR was performed in optical 96-well plates using SYBR® green in a Light Cycler 480 (Roche). Primers were designed using Primer-Blast (https://www.ncbi.nlm.nih.gov/tools/primer-blast/). The *β-tubulin2* gene (At5g05620) was used to normalize the transcript level in each sample. Primers used for RT-qPCR and stem-loop RT-qPCR are listed in Table S1. Accumulation of mature miR399f was determined by stem-loop reverse transcription quantitative PCR (Varkonyi-Gasic *et al.*, 2007). Two-way analysis of variance (ANOVA) followed by HSD (Honestly-Significant-Difference) Tukey’s test was used to analyze RT-qPCR data.

### Hormone determination

The rosettes of three-week-old WT (Col-0), miR399 OE and pho2 plants were analyzed by LC-MS for SA, SAG, JA and OPDA content as previously described in Sánchez-Bel *et al.*, 2018. Briefly, 30 mg of freeze dried material was extracted with MeOH:H2O (30:70) containing 0.01% of HCOOH containing a mixture of 10 ug. L-1 of the internal standards salicylic acid-d 5 (SA-d5) and dehydrojasmonic acid (Sigma-Aldrich). Following extraction, samples were centrifuged (15.000 rpm, 15 min) and filtered through regenerated cellulose filters. An aliquot of 20 μl was injected into a UPLC (Waters Aquity) interfaced with a Xevo TQ-S Mass Spectrometer (TQS, Waters). Hormones were quantified by contrasting with an external calibration curve of pure chemical standards of SA, SAG, JA and OPDA. Sample separation was performed with a LC Kinetex C18 analytical column of a 5 μm particle size, 2.1 100 mm (Phenomenex). Chromatographic and TQS conditions were performed as described in Sanchez-Bel *et al.* (2018). At least 6 biological replicates were analyzed per genotype and condition, each replicate consisting of leaves from at least 3 independent plants. The plant material was lyophilized prior analysis. Two-way analysis of variance (ANOVA) followed by HSD (Honestly-Significant-Difference) Tukey’s test was used to analyze data.

## ACCESSION NUMBERS

*β-Tubulin 2* (At5g62690); *PHO2* (At2g33770); *PR1* (At2g14610); *NPR1* (At1g64280); *PAD4* (At3g52430); *ERF1* (At3g23240); *PDF1.2* (At2g26020); *MYC2* (At1g32640); *VSP2* (At5g24770); *RBOHD* (At5g47910); *Ubiquitin 21*(At5g25760).

## ACKNOWLEDGEMENTS

We thank A. Molina and R.J. O’Connell for providing the *P. cucumerina* and *C. higginsianum* strains. B.V-T was a recipient of a Ph. D grant from the Ministerio de Economia, Industria y Competitividad/Agencia Estatal de Investigación/Fondo Social Europeo (BES-2016-076289). We also thank Glòria Escolà for assistance in parts of this work.

## AUTHOR CONTRIBUTIONS

BSS and T-JC devised the research project. MB, BV-T and BSS designed the experiments and analyzed the data. BV-T conducted most experiments and prepared the figures. MB-P performed infection experiments with *P. cucumerina*. HM-C collaborated in expression analyses and ROS detection. VF performed hormone analysis. BSS and BV-T wrote the article with further contributions from MB. All authors supervised and complemented the writing. BSS agrees to be responsible for contact and ensures communication.

## SUPPORTING INFORMATION

**Figure S1**. Phenotypical analysis of miR399 OE plants.

**Figure S2**. Analysis of *pho2* mutant plants under non-infection conditions.

**Figure S3**. ROS accumulation in leaves of high Pi plants (miR399 OE, *pho2* plants) under non infection conditions and infection with *P. cucumerina* by DAB staining.

**Figure S4**. ROS accumulation in leaves of high Pi plants (miR399 OE, *pho2* plants) in response to treatment with *P. cucumerina* elicitors by H_2_DCFDA staining.

**Figure S5**. ROS production in leaves of miR399 OE, *pho2* and wild type plants in response to treatment with *P. cucumerina* elicitors using the luminol assay. ROS production in wild-type plants that have been grown under different Pi supply using the luminol assay.

**Figure S6**. Expression of defense-related genes in wild type Arabidopsis plants that have been grown under different Pi supply (0.05mM, 0.1mM, or 0.25mM).

**Table S1**. List of oligonucleotides.

## REFERENCES

Aerts, N., Pereira Mendes, M. and Wees, S.C.M. Van (2021) Multiple levels of crosstalk in hormone networks regulating plant defense. Plant J., 105, 489–504.

Andersen, E., Ali, S., Byamukama, E., Yen, Y. and Nepal, M. (2018) Disease resistance mechanisms in plants. Genes (Basel)., 9, 339.

Atkinson, N.J. and Urwin, P.E. (2012) The interaction of plant biotic and abiotic stresses: From genes to the field. J. Exp. Bot., 63, 3523–3544.

Aung, K. (2006) pho2, a phosphate overaccumulator, is caused by a nonsense mutation in a microRNA399 Target Gene. Plant Physiol., 141, 1000–1011.

Ballini, E., Nguyen, T.T.T. and Morel, J.B. (2013) Diversity and genetics of nitrogen-induced susceptibility to the blast fungus in rice and wheat. Rice, 6, 1–13.

Bartel, D.P. (2004) MicroRNAs: genomics, biogenesis, mechanism, and function. Cell, 116, 281–297.

Berens, M.L., Wolinska, K.W., Spaepen, S., et al. (2019) Balancing trade-offs between biotic and abiotic stress responses through leaf age-dependent variation in stress hormone cross-talk. Proc. Natl. Acad. Sci. U. S. A., 116, 2364–2373.

Berrocal-Lobo, M., Molina, A. and Solano, R. (2002) Constitutive expression of *ETHYLENE-RESPONSE-FACTOR1* in *Arabidopsis* confers resistance to several necrotrophic fungi. Plant J., 29, 23–32.

Boccara, M., Sarazin, A., Thiébeauld, O., Jay, F., Voinnet, O., Navarro, L. and Colot, V. (2014) The Arabidopsis miR472-RDR6 silencing pathway modulates PAMP- and effector-triggered immunity through the post-transcriptional control of disease resistance Genes. PLoS Pathog., 10 (1), e1003883.

Boller, T. and Felix, G. (2009) A renaissance of elicitors: perception of microbe-associated molecular patterns and danger signals by pattern-recognition receptors. Annu. Rev. Plant Biol., 60, 379–406.

Borges, F. and Martienssen, R.A. (2015) The expanding world of small RNAs in plants. Nat. Rev. Mol. Cell Biol., 16, 727–741.

Bostock, R.M., Pye, M.F. and Roubtsova, T. V. (2014) Predisposition in plant disease: exploiting the nexus in abiotic and biotic stress perception and response. Annu. Rev. Phytopathol., 52, 517–549.

Brodersen, P., Sakvarelidze-Achard, L., Bruun-Rasmussen, M., Dunoyer, P., Yamamoto, Y.Y., Sieburth, L. and Voinnet, O. (2008) Widespread translational inhibition by plant miRNAs and siRNAs. Science (80-.)., 320, 1185–1190.

Camargo-Ramírez, R., Val-Torregrosa, B. and San Segundo, B. (2017) MiR858-mediated regulation of flavonoid-specific MYB transcription factor genes controls resistance to pathogen infection in Arabidopsis. Plant Cell Physiol., 59, 190–204.

Campos-Soriano, L., Bundó, M., Bach-Pages, M., Chiang, S.F., Chiou, T.J. and San Segundo, B. (2020) Phosphate excess increases susceptibility to pathogen infection in rice. Mol. Plant Pathol., 21, 555–570.

Castrillo, G., Teixeira, P.J.P.L., Paredes, S.H., et al. (2017) Root microbiota drive direct integration of phosphate stress and immunity. Nature, 543, 513–518.

Casacuberta, J.M., Raventós, D., Puigdoménech, P. and Segundo, B.S. (1992) Expression of the gene encoding the PR-like protein PRms in germinating maize embryos. Mol. Gen. Genet. MGG 1992 2341, 234, 97–104

Chan, C., Liao, Y.-Y. and Chiou, T.-J. (2021) The impact of phosphorus on plant immunity. Plant Cell Physiol. doi: 10.1093/pcp/pcaa168.

Chang, X. and Nick, P. (2012) Defence triggered by Flg22 and harpin is integrated into a different stilbene output in Vitis cells C.-H. Yang, ed. PLoS One, 7(7), e40446.

Chen, X. (2009) Small RNAs and their roles in plant development. Annu. Rev. Cell Dev. Biol., 25, 21–44.

Chien, P.S., Chiang, C. Bin, Wang, Z. and Chiou, T.J. (2017) MicroRNA-mediated signaling and regulation of nutrient transport and utilization. Curr. Opin. Plant Biol., 39, 73–79.

Chiou, T.J., Aung, K., Lin, S.I., Wu, C.C., Chiang, S.F. and Su, C.L. (2006) Regulation of phosphate homeostasis by MicroRNA in *Arabidopsis*. The Plant Cell, 18, 412–421.

Coolen, S., Proietti, S., Hickman, R., et al. (2016) Transcriptome dynamics of Arabidopsis during sequential biotic and abiotic stresses. Plant J., 86, 249–267.

Couto, D. and Zipfel, C. (2016) Regulation of pattern recognition receptor in plants. Nat. Rev. Immunol., 16, 537–552.

Delhaize, E. and Randall, P.J. (1995) Characterization of a phosphate-accumulator mutant of *Arabidopsis thaliana*. Plant Physiol., 107, 207–213.

Fichman, Y., Miller, G. and Mittler, R. (2019) Whole-plant live imaging of reactive oxygen species. Mol. Plant, 12, 1203–1210.

Frerigmann, H., Piotrowski, M., Lemke, R., Bednarek, P. and Schulze-Lefert, P. (2021) A Network of phosphate starvation and immune-related signaling and metabolic pathways controls the interaction between *Arabidopsis thaliana* and the beneficial fungus *Colletotrichum tofieldiae*. Mol. Plant-Microbe Interact., 34, 560–570.

Fujii, H., Chiou, T.-J., Lin, S.-I., Aung, K. and Zhu, J.-K. (2005) A miRNA involved in phosphate-starvation response in Arabidopsis. Curr. Biol., 15, 2038–2043.

Glazebrook, J. (2005) Contrasting mechanisms of defense against biotrophic and necrotrophic pathogens. Annu. Rev. Phytopathol, 43, 205–232.

Han, G.Z. (2019) Origin and evolution of the plant immune system. New Phytol., 222, 70–83.

Herrera-Vásquez, A., Salinas, P. and Holuigue, L. (2015) Salicylic acid and reactive oxygen species interplay in the transcriptional control of defense genes expression. Front. Plant Sci., 6, 1–9.

Hiruma, K., Gerlach, N., Sacristán, S., et al. (2016) Root endophyte *Colletotrichum tofieldiae* confers plant fitness benefits that are phosphate status dependent. Cell, 165, 464–474.

Huang, J., Yang, M., Lu, L. and Zhang, X. (2016) Diverse functions of small RNAs in different plantpathogen communications. Front. Microbiol., 7, 1552.

Huang, T.K., Han, C.L., Lin, S.I., et al. (2013) Identification of downstream components of ubiquitin-conjugating enzyme PHOSPHATE2 by quantitative membrane proteomics in Arabidopsis roots. Plant Cell, 25, 4044–4060.

Jagadeeswaran, G., Zheng, Y., Li, Y.F., et al. (2009) Cloning and characterization of small RNAs from Medicago truncatula reveals four novel legume-specific microRNA families. New Phytol., 184, 85–98.

Jirage, D., Tootle, T.L., Reuber, T.L., Frosts, L.N., Feys, B.J., Parker, J.E., Ausubel, F.M. and Glazebrook, J. (1999) *Arabidopsis thaliana* PAD4 encodes a lipase-like gene that is important for salicylic acid signaling. Proc. Natl. Acad. Sci. U. S. A., 96, 13583–13588.

Jones, J.D.G. and Dangl, J.L. (2006) The plant immune system. Nature, 444, 323–329.

Kadota, Y., Shirasu, K. and Zipfel, C. (2015) Regulation of the NADPH oxidase RBOHD during plant immunity. Plant Cell Physiol., 56, 1472–1480.

Keith, R.A. and Mitchell-Olds, T. (2017) Testing the optimal defense hypothesis in nature: Variation for glucosinolate profiles within plants. PLoS One, 12, e0180971.

Khan, G.A., Vogiatzaki, E., Glauser, G. and Poirier, Y. (2016) Phosphate deficiency induces the jasmonate pathway and enhances resistance to insect herbivory. Plant Physiol., 171, 632–644.

Kissoudis, C., Wiel, C. van de, Visser, R.G.F. and Linden, G. van der (2014) Enhancing crop resilience to combined abiotic and biotic stress through the dissection of physiological and molecular crosstalk. Front. Plant Sci., 5, 207.

Kraft, E., Stone, S.L., Ma, L., et al. (2016) Genome analysis and functional characterization of the E2 and RING-Type E3 ligase ubiquitination enzymes of Arabidopsis. American Society of Plant Biologists (ASPB)., 139, 1597–1611.

Lee, H.J., Park, Y.J., Kwak, K.J., Kim, D., Park, J.H., Lim, J.Y., Shin, C., Yang, K.Y. and Kang, H. (2015) MicroRNA844-guided downregulation of Cytidinephosphate Diacylglycerol Synthase3 (CDS3) mRNA affects the response of arabidopsis thaliana to bacteria and fungi. Mol. Plant-Microbe Interact., 28, 892–900.

Li, Y., Cao, X.L., Zhu, Y., et al. (2019) Osa-miR398b boosts H_2_O_2_ production and rice blast disease-resistance via multiple superoxide dismutases. New Phytol., 222, 1507–1522.

Li, Y., Zhang, Q.Q., Zhang, J., Wu, L., Qi, Y. and Zhou, J.M. (2010) Identification of microRNAs involved in pathogen-associated molecular pattern-triggered plant innate immunity. Plant Physiol., 152, 2222–2231.

Lichtenthaler, H.K. and Buschmann, C. (2001) Chlorophylls and Carotenoids: Measurement and Characterization by UV-VIS Spectroscopy. Curr. Protoc. Food Anal. Chem., 1, F4.3.1–F4.3.8.

Liu, T.-Y., Huang, T.-K., Tseng, C.-Y., Lai, Y.-S., Lin, S.-I., Lin, W.-Y., Chen, J.-W. and Chiou, T.-J. (2012) PHO2-dependent degradation of PHO1 modulates phosphate homeostasis in Arabidopsis. Plant Cell, 24, 2168–2183.

Llave, C., Xie, Z., Kasschau, K.D. and Carrington, J.C. (2002) Cleavage of Scarecrow-like mRNA targets directed by a class of Arabidopsis miRNA. Science, 297, 2053–2056.

Lorenzo, O., Chico, J.M., Sánchez-Serrano, J.J. and Solano, R. (2004) JASMONATE-INSENSITIVE1 encodes a MYC transcription factor essential to discriminate between different jasmonate-regulated defense responses in arabidopsis. Plant Cell, 16, 1938–1950.

Lu, Y.T., Li, M.Y., Cheng, K.T., Tan, C.M., Su, L.W., Lin, W.Y., Shih, H.T., Chiou, T.J. and Yang, J.Y. (2014) Transgenic plants that express the phytoplasma effector SAP11 show altered phosphate starvation and defense responses. Plant Physiol., 164, 1456–1469.

Luo, X., Li, Z., Xiao, S., Ye, Z., Nie, X., Zhang, X., Kong, J. and Zhu, L. (2021) Phosphate deficiency enhances cotton resistance to Verticillium dahliae through activating jasmonic acid biosynthesis and phenylpropanoid pathway. Plant Sci., 302, 110724.

Meldau, S., Erb, M. and Baldwin, I.T. (2012) Defence on demand: Mechanisms behind optimal defence patterns. Ann. Bot., 110, 1503–1514.

Navarro, L., Dunoyer, P., Jay, F., Arnold, B., Dharmasiri, N., Estelle, M., Voinnet, O. and Jones, J.D.G. (2006) A plant miRNA contributes to antibacterial resistance by repressing auxin signaling. Science, 312, 436–439.

Nejat, N. and Mantri, N. (2017) Plant immune system: Crosstalk between responses to biotic and abiotic stresses the missing link in understanding plant defence. Curr. Issues Mol. Biol., 23, 1–16.

Niu, D., Lii, Y.E., Chellappan, P., Lei, L., Peralta, K., Jiang, C., Guo, J., Coaker, G. and Jin, H. (2016) MiRNA863-3p sequentially targets negative immune regulator ARLPKs and positive regulator SERRATE upon bacterial infection. Nat. Commun., 7, 1–13.

Nobori, T. and Tsuda, K. (2019) The plant immune system in heterogeneous environments. Curr. Opin. Plant Biol., 50, 58–66.

O’Connell, R., Herbert, C., Sreenivasaprasad, S., Khatib, M., Esquerré-Tugayé, M.T. and Dumas, B. (2004) A novel Arabidopsis-Colletotrichum pathosystem for the molecular dissection of plant-fungal interactions. Mol. Plant-Microbe Interact., 17, 272–282.

Pandey, P., Irulappan, V., Bagavathiannan, M. V. and Senthil-Kumar, M. (2017) Impact of combined abiotic and biotic stresses on plant growth and avenues for crop improvement by exploiting physio-morphological traits. Front. Plant Sci., 8, 537.

Park, Y.J., Lee, H.J., Kwak, K.J., Lee, K., Hong, S.W. and Kang, H. (2014) MicroRNA400-guided cleavage of pentatricopeptide repeat protein mRNAs renders Arabidopsis thaliana more susceptible to pathogenic bacteria and fungi. Plant Cell Physiol., 55, 1660–1668.

Paul, S., Datta, S.K. and Datta, K. (2015) miRNA regulation of nutrient homeostasis in plants. Front. Plant Sci., 06, 232.

Pieterse, C.M.J., Does, D. Van Der, Zamioudis, C., Leon-Reyes, A. and Wees, S.C.M. Van (2012) Hormonal modulation of plant immunity. Annu. Rev. Cell Dev. Biol., 28, 489–521.

Prasch, C.M. and Sonnewald, U. (2013) Simultaneous application of heat, drought, and virus to Arabidopsis plants reveals significant shifts in signaling networks. Plant Physiol., 162, 1849–1866.

Puga, M.I., Rojas-Triana, M., Lorenzo, L. de, Leyva, A., Rubio, V. and Paz-Ares, J. (2017) Novel signals in the regulation of Pi starvation responses in plants: facts and promises. Curr. Opin. Plant Biol., 39, 40–49.

Salvador-Guirao, R., Baldrich, P., Weigel, D. and RubioSo, B.S. (2018) The microrna miR773 is involved in the arabidopsis immune response to fungal pathogens. Mol. Plant-Microbe Interact., 31, 249–259.

Sánchez-Bel, P., Sanmartín, N., Pastor, V., Mateu, D., Cerezo, M., Vidal-Albalat, A., Pastor-Fernández, J., Pozo, M.J. and Flors, V. (2018) Mycorrhizal tomato plants fine tunes the growth-defence balance upon N depleted root environments. Plant Cell Environ., 41, 406–420.

Sánchez-Vallet, A., López, G., Ramos, B., et al. (2012) Disruption of abscisic acid signaling constitutively activates Arabidopsis resistance to the necrotrophic fungus *Plectosphaerella cucumerina*. Plant Physiol., 160, 2109–2124.

Seo, J.K., Wu, J., Lii, Y., Li, Y. and Jin, H. (2013) Contribution of small RNA pathway components in plant immunity. Mol. Plant-Microbe Interact., 26, 617–625.

Shivaprasad, P. V., Chen, H.M., Patel, K., Bond, D.M., Santos, B.A.C.M. and Baulcombe, D.C. (2012) A microRNA superfamily regulates nucleotide binding site-leucine-rich repeats and other mRNAs. Plant Cell, 24, 859–874.

Smith, J.M. and Heese, A. (2014) Rapid bioassay to measure early reactive oxygen species production in Arabidopsis leave tissue in response to living *Pseudomonas syringae*. Plant Methods, 10, 6.

Snoeijers, S.S., Pérez-García, A., Joosten, M.H.A.J. and Wit, P.J.G.M. De (2000) The effect of nitrogen on disease development and gene expression in bacterial and fungal plant pathogens. Eur. J. Plant Pathol., 106, 493–506.

Song, X., Li, Y., Cao, X. and Qi, Y. (2019) MicroRNAs and their regulatory roles in plant-environment interactions. Annu. Rev. Plant Biol., 70, 489–525.

Soto-Suárez, M., Baldrich, P., Weigel, D., Rubio-Somoza, I. and San Segundo, B. (2017) The Arabidopsis miR396 mediates pathogen-associated molecular pattern-triggered immune responses against fungal pathogens. Sci. Rep., 7, 1–14.

Staiger, D., Korneli, C., Lummer, M. and Navarro, L. (2013) Emerging role for RNA-based regulation in plant immunity. New Phytol., 197, 394–404.

Tanaka, K., Choi, J., Cao, Y. and Stacey, G. (2014) Extracellular ATP acts as a damage-associated molecular pattern (DAMP) signal in plants. Front. Plant Sci., 5, 446.

Thakur, A., Verma, S., Reddy, V.P. and Sharma, D. (2019) Hypersensitive responses in plants. Agric. Rev., 40, 113–120.

Thomma, B.P.H.J., Eggermont, K., Penninckx, I.A.M.A., Mauch-Mani, B., Vogelsang, R., Cammue, B.P.A. and Broekaert, W.F. (1998) Separate jasmonate-dependent and salicylate-dependent defense-response pathways in Arabidopsis are essential for resistance to distinct microbial pathogens. Proc. Natl. Acad. Sci. U. S. A., 95, 15107–15111.

Thomma, B.P.H.J., Nürnberger, T. and Joosten, M.H.A.J. (2011) Of PAMPs and effectors: The blurred PTI-ETI dichotomy. Plant Cell, 23, 4–15.

Ton, J. and Mauch-Mani, B. (2004) β-amino-butyric acid-induced resistance against necrotrophic pathogens is based on ABA-dependent priming for callose. Plant J., 38, 119–130.

Torres, M.A., Dangl, J.L. and Jones, J.D.G. (2002) Arabidopsis gp91phox homologues Atrbohd and Atrbohf are required for accumulation of reactive oxygen intermediates in the plant defense response. Proc. Natl. Acad. Sci. U. S. A., 99, 517–522.

Torres, M.A., Jones, J.D.G. and Dangl, J.L. (2006) Reactive oxygen species signaling in response to pathogens. Plant Physiol., 141, 373–378.

Tripathi, D., Zhang, T., Koo, A.J., Stacey, G. and Tanaka, K. (2018) Extracellular ATP acts on jasmonate signaling to reinforce plant defense. Plant Physiol., 176, 511–523.

Upson, J.L., Zess, E.K., Bialas, A., Wu, C. hang and Kamoun, S. (2018) The coming of age of EvoMPMI: evolutionary molecular plant–microbe interactions across multiple timescales. Curr. Opin. Plant Biol., 44, 108–116.

Varkonyi-Gasic, E., Wu, R., Wood, M., Walton, E.F. and Hellens, R.P. (2007) Protocol: A highly sensitive RT-PCR method for detection and quantification of microRNAs. Plant Methods, 3, 1–12.

Veresoglou, S.D., Barto, E.K., Menexes, G. and Rillig, M.C. (2013) Fertilization affects severity of disease caused by fungal plant pathogens. Plant Pathol., 62, 961–969.

Versaw, W.K. and Harrison, M.J. (2002) A chloroplast phosphate transporter, PHT2;1, influences allocation of phosphate within the plant and phosphate-starvation responses. Plant Cell, 14, 1751–1766.

Wang, C., El-Shetehy, M., Shine, M.B., Yu, K., Navarre, D., Wendehenne, D., Kachroo, A. and Kachroo, P. (2014) Free radicals mediate systemic acquired resistance. Cell Rep., 7, 348–355.

Wasternack, C. and Hause, B. (2013) Jasmonates: Biosynthesis, perception, signal transduction and action in plant stress response, growth and development. An update to the 2007 review in Annals of Botany. Ann. Bot., 111, 1021–1058.

Wolinska, K.W. and Berens, M.L. (2019) Optimal Defense Theory 2.0: tissue-specific stress defense prioritization as an extra layer of complexity. Commun. Integr. Biol., 12, 91–95.

Xia, X.J., Zhou, Y.H., Shi, K., Zhou, J., Foyer, C.H. and Yu, J.Q. (2015) Interplay between reactive oxygen species and hormones in the control of plant development and stress tolerance. J. Exp. Bot., 66, 2839–2856.

Xie, Z., Allen, E., Fahlgren, N., Calamar, A., Givan, S.A. and Carrington, J.C. (2005) Expression of Arabidopsis MIRNA genes. Plant Physiol., 138, 2145–2154.

Yasuda, M., Ishikawa, A., Jikumaru, Y., et al. Antagonistic interaction between systemic acquired resistance and the abscisic acid-mediated abiotic stress response in Arabidopsis. Plant Cell, 20, 1678–1692.

Yin, H., Hong, G., Li, L., Zhang, X., Kong, Y., Sun, Z., Li, J., Chen, J. and He, Y. (2019) MiR156/SPL9 regulates reactive oxygen species accumulation and immune response in Arabidopsis thaliana. Phytopathology, 109, 632–642.

Zhang, L., Zhang, F., Melotto, M., Yao, J. and He, S.Y. (2017) Jasmonate signaling and manipulation by pathogens and insects. J. Exp. Bot., 68, 1371–1385.

Zhao, H., Sun, R., Albrecht, U., et al. (2013) Small RNA profiling reveals phosphorus deficiency as a contributing factor in symptom expression for citrus huanglongbing disease. Mol. Plant, 6, 301–310.

Zheng, X.Y., Spivey, N.W., Zeng, W., Liu, P.P., Fu, Z.Q., Klessig, D.F., He, S.Y. and Dong, X. (2012) Coronatine promotes pseudomonas syringae virulence in plants by activating a signaling cascade that inhibits salicylic acid accumulation. Cell Host Microbe, 11, 587–596.

