## supplemental figures and table for "Phosphate-induced resistance to pathogen infection in Arabidopsis"

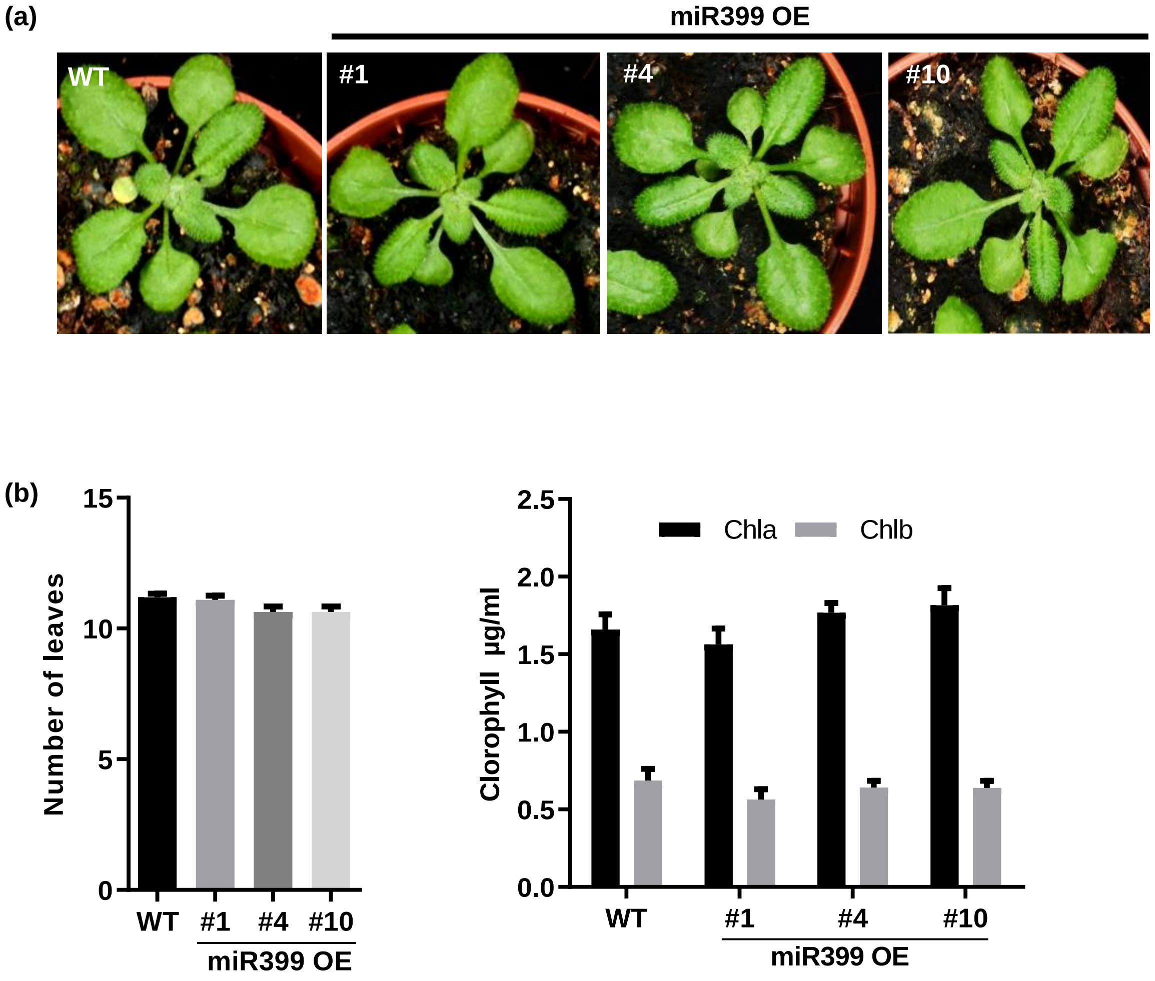
(a)
miR399 OE
WT
#1
#4
#10
(b)
15
10
5
0
Number of leaves
WT
#1
#4
#10
miR399 OE
2.5
2.0
1.5
1.0
0.5
0.0
Clorophyll μg/ml
Chla
Chlb
WT
#1
#4
#10
miR399 OE

### Figure S1

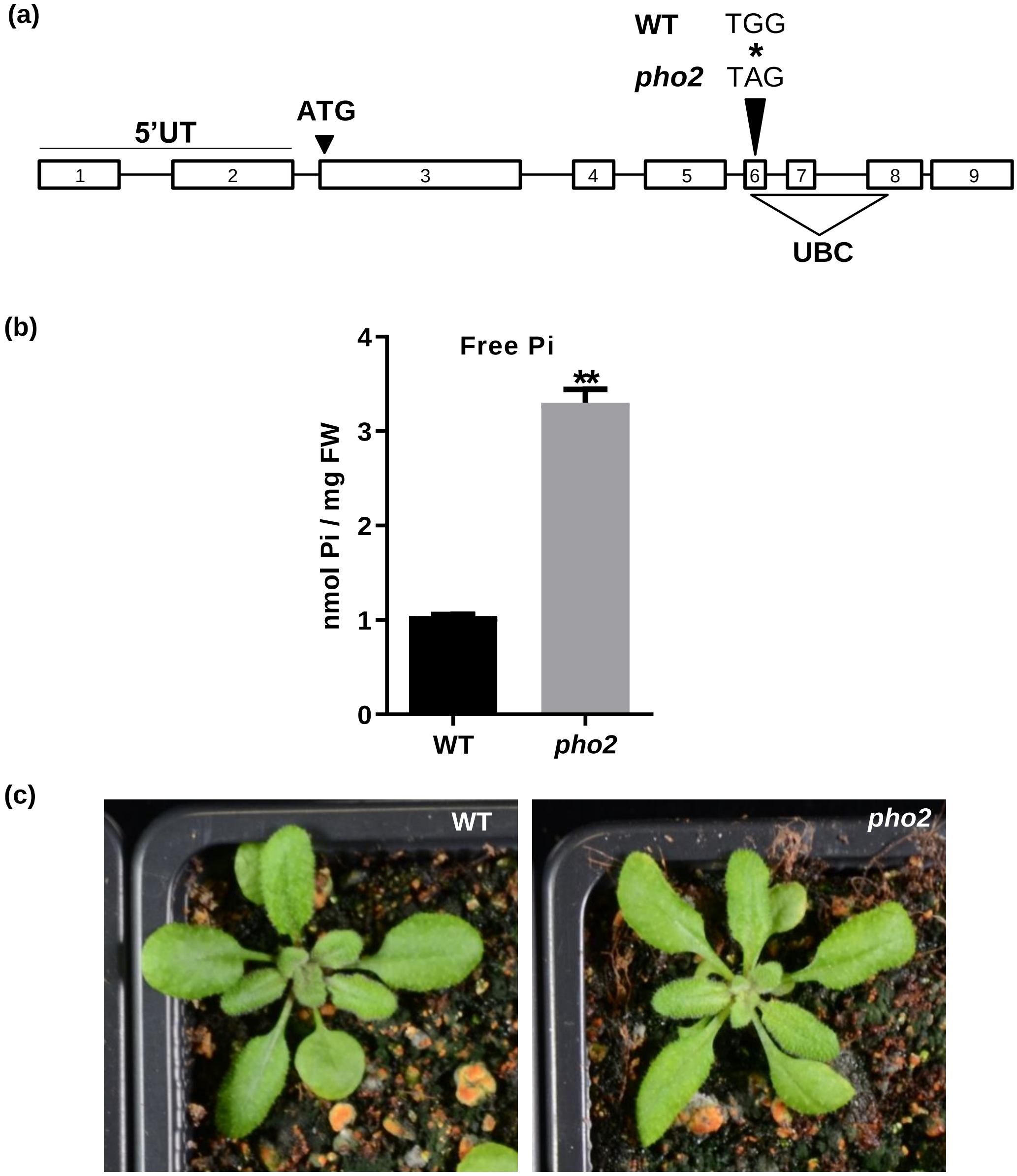
(a)
WT TGG
*
pho2 TAG
ATG
5'UT
1
2
3
4
5
6
7
8
9
UBC
(b)
Free Pi
**
nmol Pi / mg FW
0
1
2
3
4
WT
pho2
(c)
WT
pho2

### Figure S2

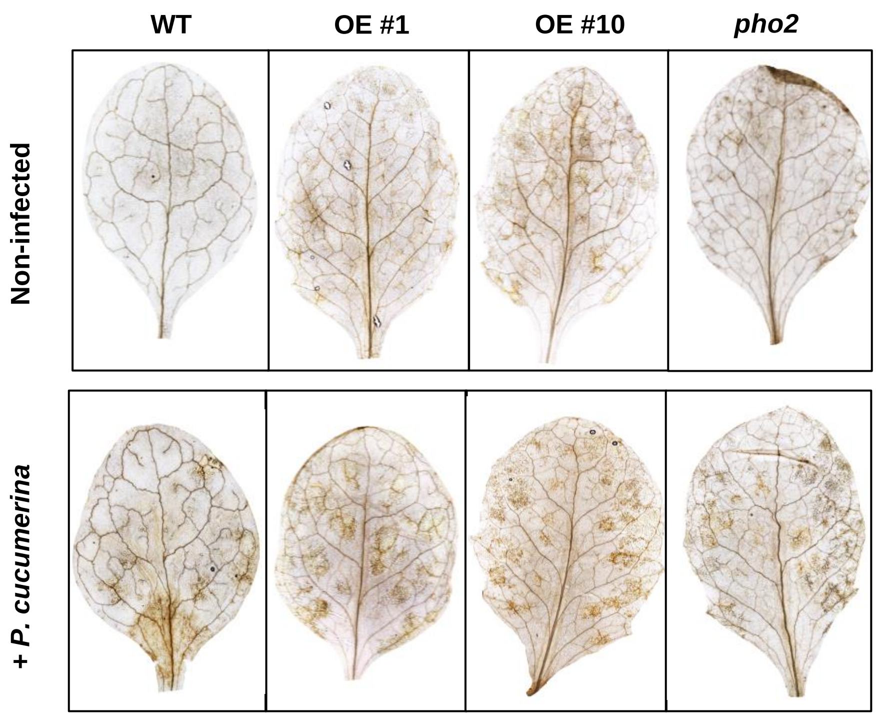


**Figure S3**

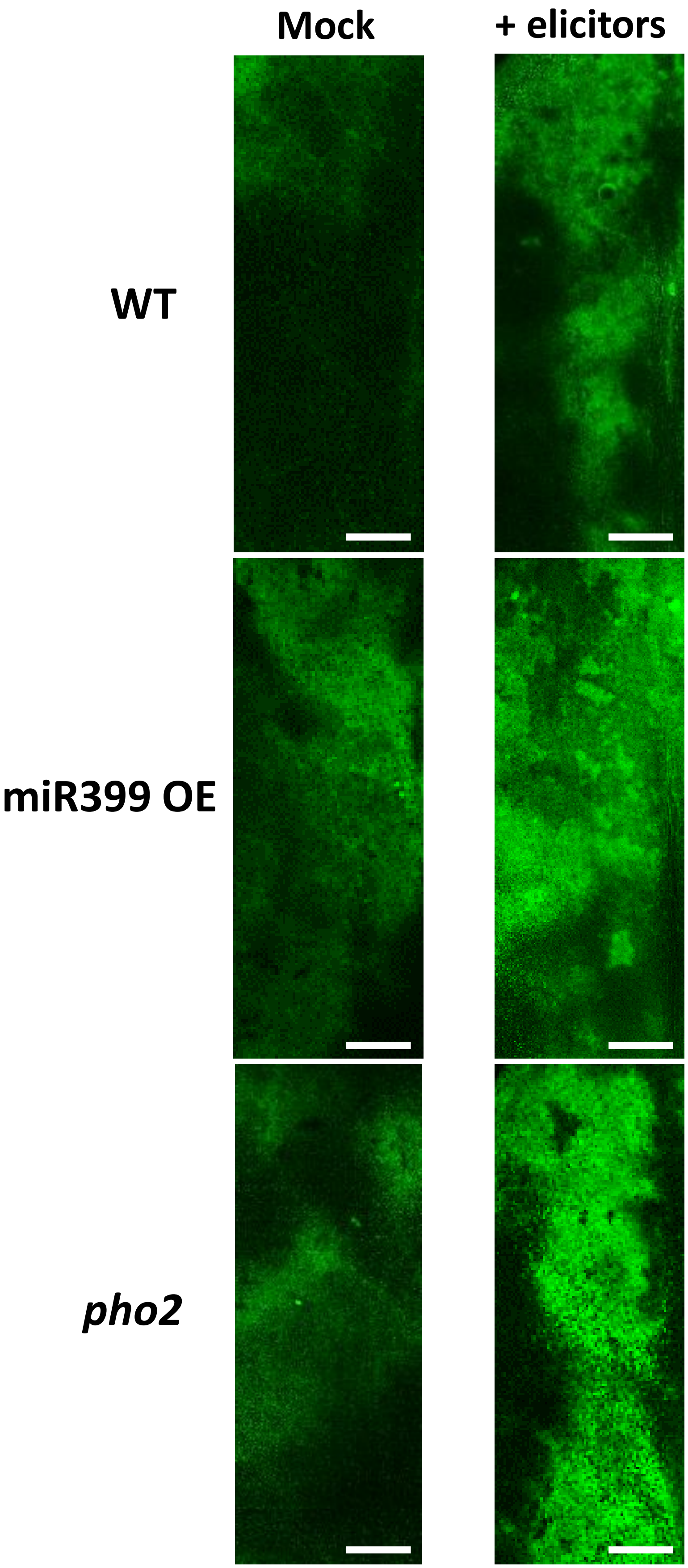


Figure S4

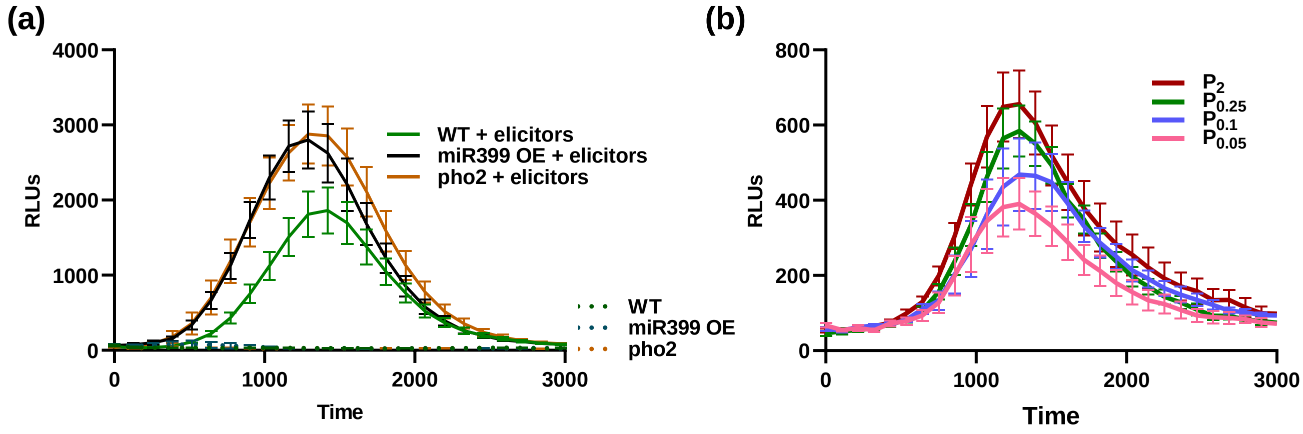
(a)
4000
3000
2000
1000
0
RLUs
0
1000
2000
3000
Time
WT + elicitors
miR399 OE + elicitors
pho2 + elicitors
WT
miR399 OE
pho2
(b)
800
600
400
200
0
RLUs
0
1000
2000
3000
Time
P2
P0.25
P0.1
P0.05

**Figure S5**

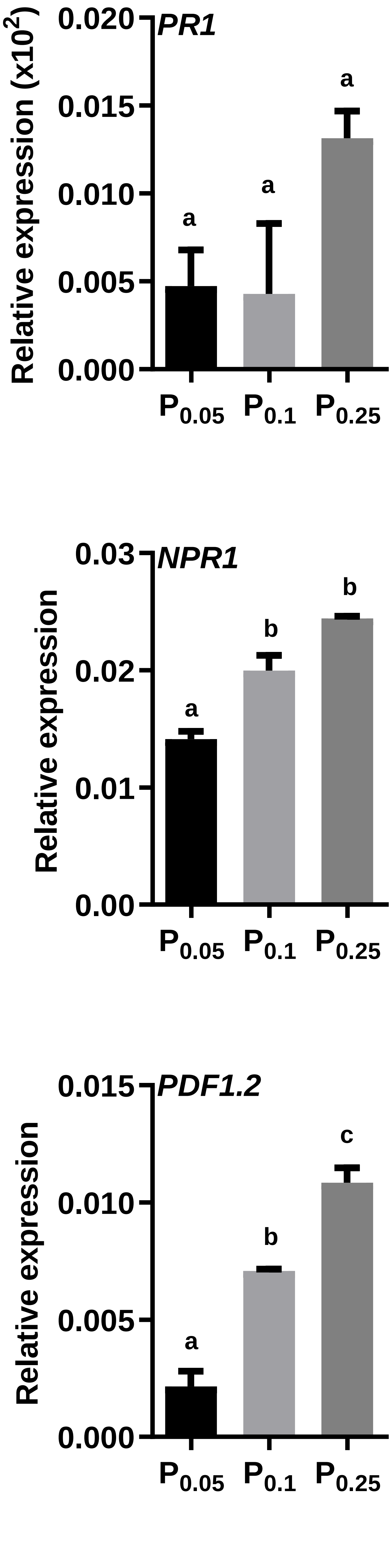

PR1
Relative expression (x10^2)
0.020
0.015
0.010
0.005
0.000
a
a
a
P0.05
P0.1
P0.25
NPR1
Relative expression
0.03
0.02
0.01
0.00
a
b
b
P0.05
P0.1
P0.25
PDF1.2
Relative expression
0.015
0.010
0.005
0.000
a
b
c
P0.05
P0.1
P0.25


**Figure S6**

Table S1. List of oligonucleotides

| Gene name | Accession Number | | Sequence (5'-3') |
|---|---|---|---|
| **For expression analysis** | | | |
| *β-Tubulin 2* | *At5g62690* | Fw | TGTTCAGGCGAGTGAGTGAG |
| | | Rv | ATGTTGCTCTCCGCTTCTGT |
| *Pre-miR399* | | Fw | TGCATAAATGTTTGTGGTGAGC |
| | | Rv | GAATTACCGGGCAAATCTCCT |
| *Pre-miR827* | | Fw | ACATGTTGATCATCCTTGTGTTGA |
| | | Rv | CCAAGAAGCGATGCAAAACCA |
| *miR399 stem loop* | | Rt | GTTGGCTCTGGTGCAGGGTCCGAGGTATTCGCACCAGAGCCAACCCGGGC |
| | | Fw | GCGGTGCCAAAGGAGATTT |
| *PHO2* | *At2g33770* | Fw | AGGTTTGAAGCTCCACCCTCA |
| | | Rv | CCCAAGATGTGATTGGAGTTCC |
| *PR1* | *At2g14610* | Fw | GATGTGCCAAAGTGAGGTGTAA |
| | | Rv | GGCTTCTCGTTCACATAATTCC |
| *NPR1* | *At1g64280* | Fw | CCGGAAGAGCTTGTTAAAGAGA |
| | | Rv | ATCCGAGTCAAGTGCCTTATGT |
| *PAD4* | *At3g52430* | Fw | CGAATACATTGGTGACGAAGAA |
| | | Rv | ACCCATTTTGCACTTGAACTCT |
| *ERF1* | *At3g23240* | Fw | AACACTCGATGAGACGGAGAAT |
| | | Rv | CTCCCAAATCCTCAAAGACAAC |
| *PDF1.2* | *At2g26020* | Fw | CAACAATGGTGGAAGCACAG |
| | | Rv | CTTGCATGCATTGCTGTTTC |
| *MYC2* | *At1g32640* | Fw | ATTAATGACCCGATTGGAACAC |
| | | Rv | TGAGCTACCGTTCTCAAACTGA |
| *VSP2* | *At5g24770* | Fw | CTCGTCGATTCGAAAACCAT |
| | | Rv | TTCTGCAGTTGGCGTAGTTG |
| *RBOHD* | *At5g47910* | Fw | ATTACAAGCACCAAACCAG |
| | | Rv | TTCTCCGACCATCTCACTA |
| **For fungal DNA quantification** | | | |
| *P.cucumerina_tubulin* | | Fw | CAAGTATGTTCCCCGAGCCGT |
| | | Rv | GAAGAGCTGACCGAAGGGACC |
| *Ch_ITS2* | | Fw | AAAGGTAGTGGCGGACCCTC |
| | | Rv | GGCAAGAGTCCCTCCGGAt |
| *Ubiquitin21* | *At5g25760* | Fw | AAAGGACCTTCGGAGACTCCTTACG |
| | | Rv | GGTCAAGAATCGAACTTGAGGAGGTT |
